# Chemotaxis of branched cells in complex environments

**DOI:** 10.1101/2025.05.27.656457

**Authors:** Jiayi Liu, Jonathan E. Ron, Giulia Rinaldi, Ivanna Williantarra, Antonios Georgantzoglou, Ingrid de Vries, Michael Sixt, Milka Sarris, Nir S. Gov

## Abstract

Cell migration *in vivo* is often guided by chemical signals. Such chemotaxis, such as performed by immune cells migrating to a wound site, is complicated by the complex geometry inside living tissues. In this study, we extend our theoretical model of branched-cell migration on a network by introducing chemokine sources to explore the cellular response. The model predicts a speed-accuracy tradeoff, whereby slow cells are significantly more accurate and able to follow efficiently a weak chemoattractant signal. We then compare the model’s predictions with experimental observations of neutrophils migrating to the site of laser-inflicted wound in a zebrafish larva fin, and migrating *in-vitro* inside a regular lattice of pillars. We find that the model captures the details of the sub-cellular response to the chemokine gradient, as well as the large-scale migration response. This comparison suggests that the neutrophils behave as fast cells, compromising their chemotaxis accuracy, which explains the functionality of these immune cells.

## Introduction

Tissues have complex geometries and present a great diversity of topologies which force embedded motile cells to deform and exhibit a plethora of branched shapes [1]. Archetypal examples include immune cells that continuously scan the tissues in search for signals and need to migrate within the complex tissues towards sites of wound and infection [2–4] (Fig. 1), or cancer cells that spread from the main tumor [5]. In order to efficiently migrate towards their target, branched cells must coordinate their branch dynamics to navigate the microenvironment and decide on the new migration direction, in a process termed directional decision-making (DDM) [6, 7]. Despite its clear biological importance, the mechanisms of branched cell migration and chemotaxis through complex environments remain poorly understood [8]. Furthermore, there is currently a lack of theoretical framework for tackling these questions.

**Fig. 1:**
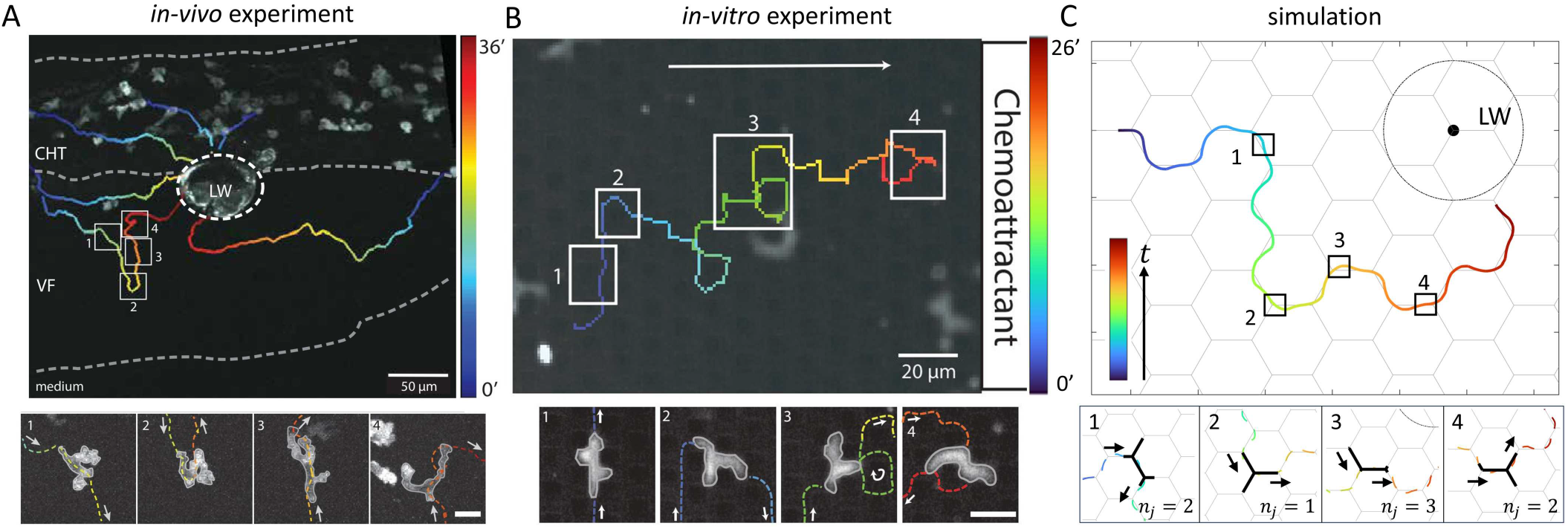
(A) Morphological dynamics of neutrophil swarming during migration from the caudal hematopoietic tissue (CHT) toward a laser-induced wound (LW) at the ventral fin (VF)–CHT boundary in transgenic zebrafish larva expressing the calcium indicator GCamp6f (see SI section S-1 for more details). Top panel: Representative trajectory plots of neutrophils migrating toward the LW, with trajectory color denoting migration time. White boxes highlight time points along the trajectory of an individual cell, corresponding to the images in the bottom panels (1–4). White arrows indicate the direction of migration. Scale bars: 50 *µ*m for the trajectory plots and 10 *µ*m for the enlarged images. (B) Trajectory and snapshots of the cell shape, while performing chemotaxis inside a regular lattice of pillars (see SI section S-2 for more details) [9]. (C) Simulation of a branched cell moving toward a chemokine point source. Top panel: A representative trajectory of the cell’s center of mass (C.O.M.) during migration, with trajectory color denoting migration time. Black boxes highlight time points along the trajectory of the cell, corresponding to the images in the bottom panels (1–4). Black arrows indicate the direction of migration. Key parameters: *C/c*_0_ = 0.01, *ϵ* = 0.2, *d* = 3, *β*_0_ = 8, σ = 0.8.

We have recently developed a theoretical model describing the shape dynamics of branched migrating cells performing DDM over single junctions [10], and the spontaneous polarization and migration of highly branched cells on hexagonal networks [11]. The coarse-grained model describes cells migrating on an hexagonal network, composed of linear segments (Fig. 1C), with an internal mechanism for spontaneous cellular polariaztion and migration.

Here we utilize this theoretical model to explore the chemotaxis characteristics of branched cells. To test the simulations we compared them to two systems (Fig. 1), one *in vivo* using tissue injury in zebrafish [4, 12] and one *in-vitro* where cells migrate in a regular lattice of pillars [9]. We validate the model by comparing to experimental data of the shape and migration dynamics of neutrophils as they respond to a signal that recruits them to a wound, in the zebra-fish skin [4]. Our model provides insights into the constraints that determine the optimal migration properties of neutrophils.

### Chemotaxis over a single junction

We start by studying the chemotaxis of a cell when migrating over a single junction (Fig. 2A). We extend our previous model [10, 13], by introducing the effect of an external chemical signal (see Supplementary Information section S-3 for the model equations). The binding of the chemokine to the receptors at the leading edge of the cell triggers a local enhancement of the activity of the actin polymerization machinery [14]. This is represented in our model by an increase in the value of the parameter *β*, which describes the strength of the actin flow and protrusive force acting at the tips of the arms of the branched cell. For a single junction we simplify the signal by assuming that it is localized along one of the arms that extend from the junction (arm 3, Fig. 2A), described as:

**Fig. 2:**
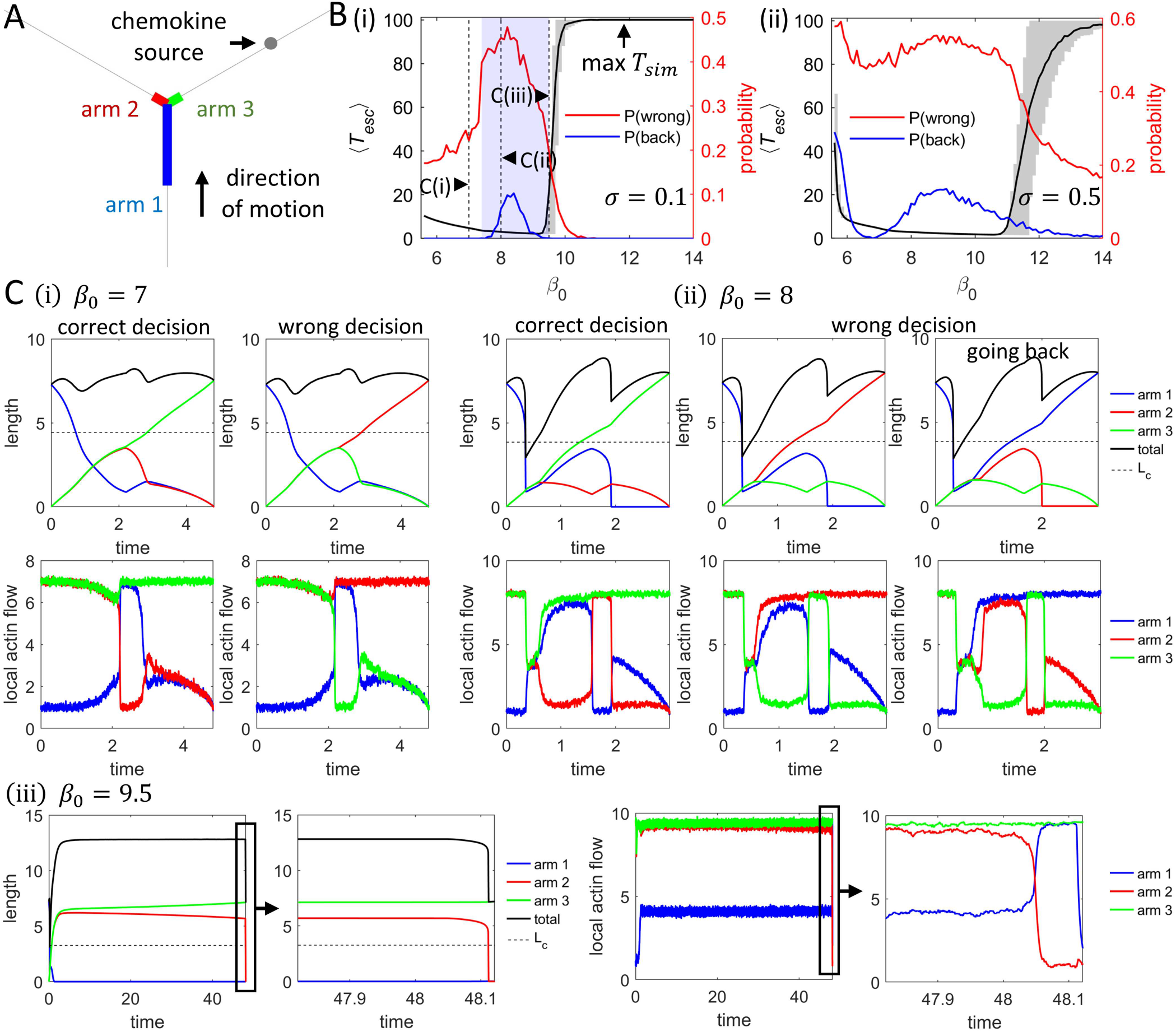
Chemotaxis dynamics over a single-junction with a localized chemokine source. (A) Schematic representation of the model. (B) Mean time for the cell to traverse the junction *T*_*esc*_ (black), mean error rate *P* (*wrong*) (the probability of leaving along arm 2, red), and the mean probability of being “reflected” at the junction *P* (*back*) (leaving along arm 1, blue), as functions of *β*_0_. Maximum simulation time: *T* = 100. The blue shaded region in (i) marks the range where cells undergo stick-slip events, and a cell can reverse its path (*P* (*back*) *>* 0). Panels (i) and (ii) correspond to two values of the internal noise in the actin polymerization activity σ = 0.1 and σ = 0.5, respectively. (C) Time series of length of the arms and total cell length (top), as well as local actin flows at the tips of arms (bottom), for representative events: making the correct decision (leaving along arm 3), making the wrong decision while moving forward (leaving along arm 2), and making the wrong decision by turning back (leaving along arm 1). Panels (i), (ii), and (iii) correspond to *β*_0_ = 7.0, *β*_0_ = 8.0 and *β*_0_ = 9.5, respectively. Other key parameter: *ϵ* = 0.001.

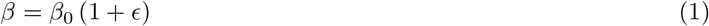

where *β*_0_ represents the actin polymerization speed parameter in the absence of external signals, and *ϵ* is the increase factor due to the chemokine signal. For the tips of the other two arms (arms 1 and 2), the polymerization speed parameter remains unchanged, with *β* = *β*_0_.

As in our previous work [10], the model equations (see SI section S-3) are normalized by the timescale of the inverse of the focal adhesion disassembly rate (5–30 min [15–18]) and by the length scale of the rest length of the cell (10–100 *µm* on one-dimensional tracks [13]). In the rest of this paper, we kept all the parameters of the cells as were calibrated to fit the observed dynamics of typical motile cells [10] (Table. S-1).

Our main free control parameter is the value of the actin polymerization (and retrograde flow) activity, *β*_0_. This parameter has a lower bound given by the minimal value (*β*_*c*_) that allows the cells to polarize [13] and migrate over the network [10]. This value is weakly dependent on the cell shape, increasing with the number of network nodes and branches that the cell has. To ensure that our model cell is able to migrate, we therefore explore the dynamics for *β*_0_ *> β*_*c*_ (where *β*_*c*_ 5 − 6 for the model parameters chosen here [10]). Above a higher critical value,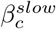the model predicts that the cell may get stuck in a “slow-mode” process, where the two leading arms are highly elongated while the back is stuck at the same node, until the competition between the two elongated arms is resolved [10]. We find that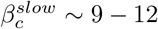for the model parameters chosen here (Table. S-1). These considerations determine the range of *β*_0_ values that we explore in this paper (see Fig. 2B for example).

In Fig. 2B(i,ii) we plotted for a fixed small bias *ϵ* = 0.001 (Eq.1) and two different levels of the noise the mean escape time that it takes for the cell to migrate over the junction, ⟨*T*_*esc*_⟩, and the probability of making the wrong decision, *P* (*wrong*), as functions of *β*_0_ (Fig. 2B). The wrong decision indicates that the cell does not leave the junction along the path that leads to the chemokine source (arm 3), but rather leaves along either arm 2 or arm 1. The proportion of events where the cell is “reflected” by the junction, i.e. turn back and migrate along arm 1, are denoted by the blue line. The region where stick-slip migration over the junction occurs is denoted by the blue shading, and corresponds to the region where the reflection probability is nonzero. Typical examples of the cellular dynamics during different cases of directional decision-making are shown in Fig. 2C (for two values of *β*_0_ denoted by vertical dashed lines in Fig. 2B(i)).

For weak noise (σ = 0.1, Fig. 2B(i))), the mean escape time (⟨*T*_*esc*_⟩) decreases as *β*_0_ increases, which is expected as the cell velocity increases with an increase of actin polymerization activity [10]. At large values of *β*_0_ ∼ 10 the mean escape time sharply increases, as the cell tends to remain for a long period of time in the “slow-mode”, with two highly elongated arms that are locked in symmetric competition (see Fig. 2C(iii)), consistent with the cell behavior without chemotaxis [10]. We see that in this mode the rear arm of the cell (arm 1) shrinks approximately to zero and the two new arms remain elongated with similar lengths for a long time, until one of the arms rapidly retracts.

The accuracy, as quantified by the mean error probability *P* (*wrong*), shows an opposite trend, exhibiting a “speed-accuracy tradeoff”. Specifically, cells that spend less time over the junction are more likely to make incorrect decisions and miss their way to the target. The abrupt increase in *P* (*wrong*) for small noise at *β*_0_ ≈ 8 arises from the emergence of a finite probability for the cell to reflect along the original path (blue line in Fig. 2B(i)). This behavior occurs exclusively during stick-slip events [10, 13], which is observed only for larger *β* (in the blue-shaded regime of Fig. 2B(i)). Examples of smooth and stick-slip migrations are shown in Fig. (2C(i,ii)). For smooth motion, the only possible incorrect decision is to leave the junction along the new path without the source (arm 2). However, since during stick-slip events the cell length decreases below the critical polarization length *L*_*c*_ (denoted by the horizontal dashed line in Fig. 2C(i,ii)), cells may also escape the junction by returning along their original path (arm 1). The duration that the cell spends with a length below *L*_*c*_, and loses its polarity, causes the sharp increase in accuracy errors (similar to behavior at high noise and very low *β*_0_, as discussed below).

As *β*_0_ increases further, the cell transitions into the “slow mode”, characterized by two elongated and competing arms along the new paths, as observed in the one-junction model [10] and shown in Fig. 2C(iii), for two noise levels. We calculated the probability of slow mode occurrence, *P* (*slow*), as a function of *β*_0_ (Fig. S-3A), and found that it closely aligns with the trend of the mean escape time (⟨*T*_*esc*_⟩) versus against the actin polymerization speed baseline (*β*_0_) (Fig. 2B(i)). Additionally, we computed the probability of making incorrect decisions in the fast and slow modes (red solid and dashed lines in Fig. S3-A, respectively), showing that the slow mode exhibits significantly higher accuracy. Finally, for specific values of *β*_0_ (denoted by black dashed lines in Fig. S-3A), we plotted in Fig. S-3B(i-iii) the distribution of escape times *T*_*esc*_ (histograms) and the value of *P* (*wrong*) as function of *T*_*esc*_ (black line). We observe a complex relation between the speed of individual events and their accuracy. This complex relation arises from two competing effects: slower events can be more accurate due to the speed-accuracy relation that appears on the average values. On the other hand, higher escape times can be associated with events of polarity loss, which give rise to higher error rates.

For larger noise (σ = 0.5), we find a similar tradeoff as for the low noise, except at very low values of *β*_0_ (Fig. 2B(ii)). In this limit, due to the large noise the cell loses its polarization when migrating over the junction, and has an almost equal probability of leaving along any of the three arms, leading to a *P* (*wrong*) ∼ 2*/*3.

Taken together, these results confirm the existence of a tradeoff between speed and accuracy of cellular DDM during chemotaxis. In the next section we explore the consequences of this behavior for the large-scale chemotactic migration of cells over a hexagonal network.

### Branched cell chemotaxis on a hexagonal network

We next studied the migration dynamics of a branched cell on a hexagonal network, in the presence of a global chemokine gradient.

Our treatment of a branched cell that spans more than one junction was described in [11], and explained in more details in SI section S-3. Here, we expose these branched cells to a chemokine concentration field that has an exponential form:

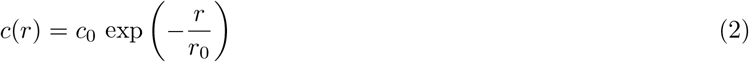

where *c*_0_ is the chemokine concentration at the source position, *r*_0_ is the decay length of the chemokine concentration, and *r* is the distance from the tips of the cellular arms to the chemokine source. The local actin polymerization activity at each arm tip is enhanced due to the chemokine concentration profile given by (similar to Eq.1):

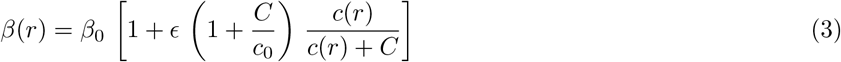

where *β*_0_ is the actin polymerization speed in the absence of external signals, *ϵ* is the maximum enhancement of *β*, and *C* is the saturation concentration for detecting the signal. The dimensionless factor (1 + *C/c*_0_) normalizes the enhancement to be maximal at *r* = 0.

In this study, we model a chemokine source along a one-dimensional line located at *y*_*source*_ = 8*d*, where *d* is the edge length of the hexagons in the network. The cell’s arrival is defined as the point where any of its arm tips reaches *y*_*arr*_ = 6.5*d* (black dash-dotted line in the sketch in Fig. 3A). The chemokine concentration is therefore written as:

**Fig. 3:**
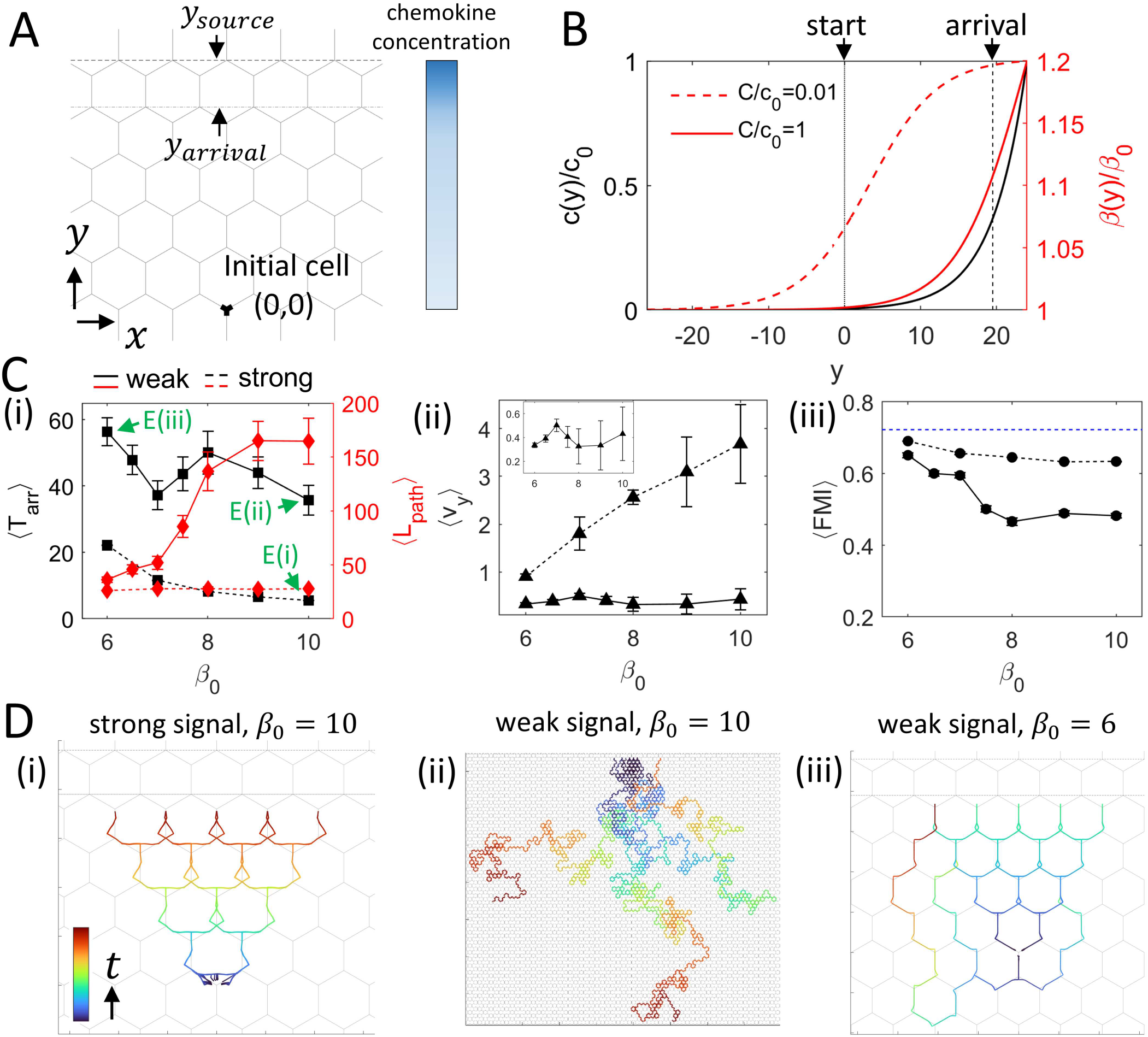
Multiple-junction model with a chemokine line source. (A) Schematic representation of the model. The gray dashed line indicates the position of the chemokine source, while the gray dash-dotted line marks the arrival position. (B) Normalized chemokine concentration, *c*(*y*)*/c*_0_ (black line), and the enhancement of actin polymerization speed, *β*(*y*)*/β*_0_, as functions of *y* in the strong signal regime (red dashed line) and the weak signal regime (red solid line). (C) (i) ⟨*T*_*arr*_⟩ and ⟨*L*_*path*_⟩ as functions of *β*_0_. (ii) ⟨*v*_*y*_ ⟩ as a function of *β*_0_. The inset corresponds to the weak signal regime. (iii) Forward Migration Index ⟨*FMI*⟩ (black solid line) and maximal theoretical *FMI* (blue dashed line) as functions of *β*_0_. Black solid lines and dashed lines correspond to the weak and strong signal regimes in (B), respectively. (D) Typical trajectories of the cell’s C.O.M. during a simulation with (i) strong signal, *β*_0_ = 10.0, (ii) weak signal, *β*_0_ = 10.0, and (iii) weak signal, *β*_0_ = 6.0. Maximal simulation time: *T* = 1000. Other key parameters: *ϵ* = 0.2, *d* = 3, σ = 0.1.

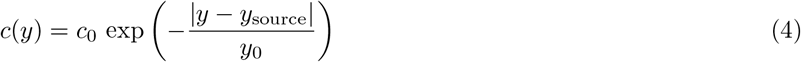

where *y* is the *y*-coordinate of the arm tip.

We fixed the chemokine profile parameters (*ϵ* = 0.2 and *y*_0_ = 1.5*d*) and vary the parameter *C/c*_0_ which represents different saturation regimes of the cells in the chemokine gradient. For *C/c*_0_ = 0.01, the chemokine source attracts the cell even when it is far away (Fig. 3B), representing the strong signal (high saturation) regime. For *C/c*_0_ = 1, the chemokine gradient is only effective when the cell is very close to the source (Fig. 3B), representing the weak signal (low saturation) regime. At the start of each simulation, the cell is symmetrically positioned at the origin (*x, y*) = (0, 0) and allowed to spread, polarize and migrate. The simulation ends when the cell reaches the source, or when the maximal simulation time is reached *T*_*max*_ = 1000.

We evaluated the efficiency of the chemotactic migration by plotting the mean arrival time, *T*_*arr*_, which reflects both the speed and accuracy of the DDM during the migration (Fig. 3C(i)). Additionally, we calculated the mean distance of the cell’s center-of-mass (C.O.M.) trajectory, *L*_path_ (Fig. 3C(i), mean speed towards the chemokine source (⟨*v*_*y*_⟩, Fig. 3C(ii)) and forward migration index (FMI, Fig. 3C(iii)).

In the strong signal regime, the cells rarely take wrong turns and migrate along directed paths to the chemokine source (see typical trajectories for *β*_0_ = 10 in Fig. 3D(i)). As a result, ⟨*L*_*path*_⟩ is small and independent of *β*_0_ (Fig. 3C(i), red line). The high accuracy is also reflected in ⟨*v*_*y*_⟩ (Fig. 3C(ii)), which is increasing almost linearly with *β*_0_. Similarly, ⟨*T*_*arr*_⟩ decreases monotonically as *β*_0_ increases (Fig. 3C(i), black dashed line), since the cell’s speed increases with *β*_0_, without reducing its accuracy.

In the weak signal regime, we find a more complex behavior as a function of increasing *β*_0_. At the lowest values of *β*_0_, we find that ⟨*L*_*path*_⟩ and FMI are very similar although ⟨*T*_*arr*_⟩ is longer for strong signals (Fig. 3), indicating that these slowest cells are very accurate in their chemotaxis migration even in response to a very weak signal.

As *β*_0_ increases, the ⟨*L*_*path*_⟩ increases monotonically (Fig. 3C(i), red solid line), indicating a higher probability of making incorrect DDM, as turning away from the chemokine source. This fits with the speed-accuracy relation we found on a single junction in Fig. 2B. It is most clearly shown by plotting typical trajectories of the cell’s C.O.M. at large *β*_0_ = 10 (Fig. 3D(ii)).

At intermediate values, around *β*_0_, we find a significant minimum of ⟨*T*_*arr*_⟩ (Fig. 3C(i), black solid line). This is due to the slower increase in the error rate (quantified by the FMI in Fig. 3C(iii)) in comparison to the increase in speed, with an increase of *β*_0_ (Fig. 3C(ii)). At higher *β*_0_ values, the error rate increases faster than the speed, which increases ⟨*T*_*arr*_⟩ that reaches a peak around *β*_0_ ∼ 8 (Fig. 3C(i), black solid line). The non-monotonous behavior of ⟨*T*_*arr*_⟩ that we find, largely follows the speed-accuracy trends that we found on a single junction (Fig. 2B(i)).

However, despite the wrong turns, the higher overall speed for the high values of *β*_0_ ∼ 10 results in a low ⟨*T*_*arr*_⟩ (Fig. 3C(i), black solid line), which is also reflected by ⟨*v*_*y*_⟩ (Fig. 3C(ii), solid line). Note that for the very small fraction of cells (∼ 1 − 2%) that do not arrive at the target within the *T*_max_, we assigned *T*_arr_ = 1000.

In the SI section S-4, we investigated the effect of cellular internal noise on chemotactic migration in both the strong and weak signal regimes (Fig. S-4A,B). We also investigated the migration dynamics of cells on large grids (*d* = 7.5) (Fig. S-4C,D). In this case, we found that cells with large *β*_0_ values occasionally exhibit slow-mode events, similar to the behavior observed in the one-junction case (Fig. 2C(iii)). We show that the slow-mode appears along the trajectory, which causes the mean arrival time to increase and the FMI (accuracy) to improve in the strong-signal regime (see Fig. S-4D,F). Note that a similar change in the migration dynamics can be induced by keeping the grid size fixed while the cell length is modified, for example by changing the cell’s contractility and stiffness parameter *k*. The slow-mode events are also shown in simulations towards a point-like chemokine source (Figs.S-9,5), and have two types: either the two long arms form in the direction of migration (Fig.5) as in a single junction (Fig. 2C(iii)), or one of the arms form along the direction from which the cell arrived at the junction, following a stick-slip event (Fig.S-9). We also give additional detailed analysis of the distribution of arrival times, and how they are related to the errors that the cell performs along its migration path (SI section S-5, Fig.S-5).

From the theoretical studies above, we can summarize the following conclusions regarding the best strategies that an immune cell may adopt in order to arrive to a site of a wound or infection using chemotaxis:

- In the presence of a strong signal, as may be expected close to the target site (wound or infection), cells with higher actin polymerization activity *β*_0_ arrive faster, and therefore can prevent bacterial entry most effectively (Fig. 3C(i)).
- In the presence of a weak signal, far from the target site, we find that the slowest cells are able to maintain accurate and directed chemotaxis migration paths. Despite the high accuracy, the arrival time for these slow-moving cells is relatively long.
- When the signal is weak, the fastest arrival times are found for a narrow range of intermediate cellular activities (*β*_0_ ∼ 7) (Fig. 3D(i)). In this intermediate regime, the chemotaxis is most efficient. This is demonstrated by short arrival times and low energy expenditure (which may allow them to have a longer lifespan or remain more potent, for example).
- For weak signals far from the target site, cells with high *β*_0_ exhibit a high percentage of meandering paths due to lower chemotactic accuracy (compared to slower cells). Nevertheless, they can have a relatively short mean arrival time, due to their high speeds. On grids of larger size these highly active cells may get further slowed down due to getting transiently trapped in slow-mode events (Figs. S-3,S-8,5). Efficient recruitment of these high-activity cells from large distances may therefore not rely only on the initial signal from the wound/infection site, but utilize further strategies to enhance the chemical guidance signal.

Comparison of the model with the chemotaxis of neutrophils

Next, we compare our theoretical model with *in-vivo* experimental data of neutrophils migrating towards laser-inflicted wounds (LW) [12] in the fin of a zebrafish larva [4].

To simplify the analysis of the chemotaxis within our theoretical model, we assume a spatially linear variation of the chemokine concentration (Fig.S-6A), increasing along the *y*-axis (for more details of the choice of parameters for the concentration profile see SI section S-6).

In Fig. 4A, we show an example of the dynamics of a single migrating neutrophil, which initially moves away from the LW (the LW is located at the +*y*-direction). The cell is observed to respond to the chemical signal from the LW by slowing down, pausing, and rotating towards the LW. The dynamic of the cell’s speed along the *y*-direction, and its angle with respect to the *y*-direction, during this process are shown in Fig.S-5C. While migrating towards the LW, the neutrophil displays large speeds and large directional variability (see time*>* 6 min in Fig.S-6C). Further properties of the simulated cell’s response to the chemokine gradient, are discussed in SI section S-6.

**Fig. 4:**
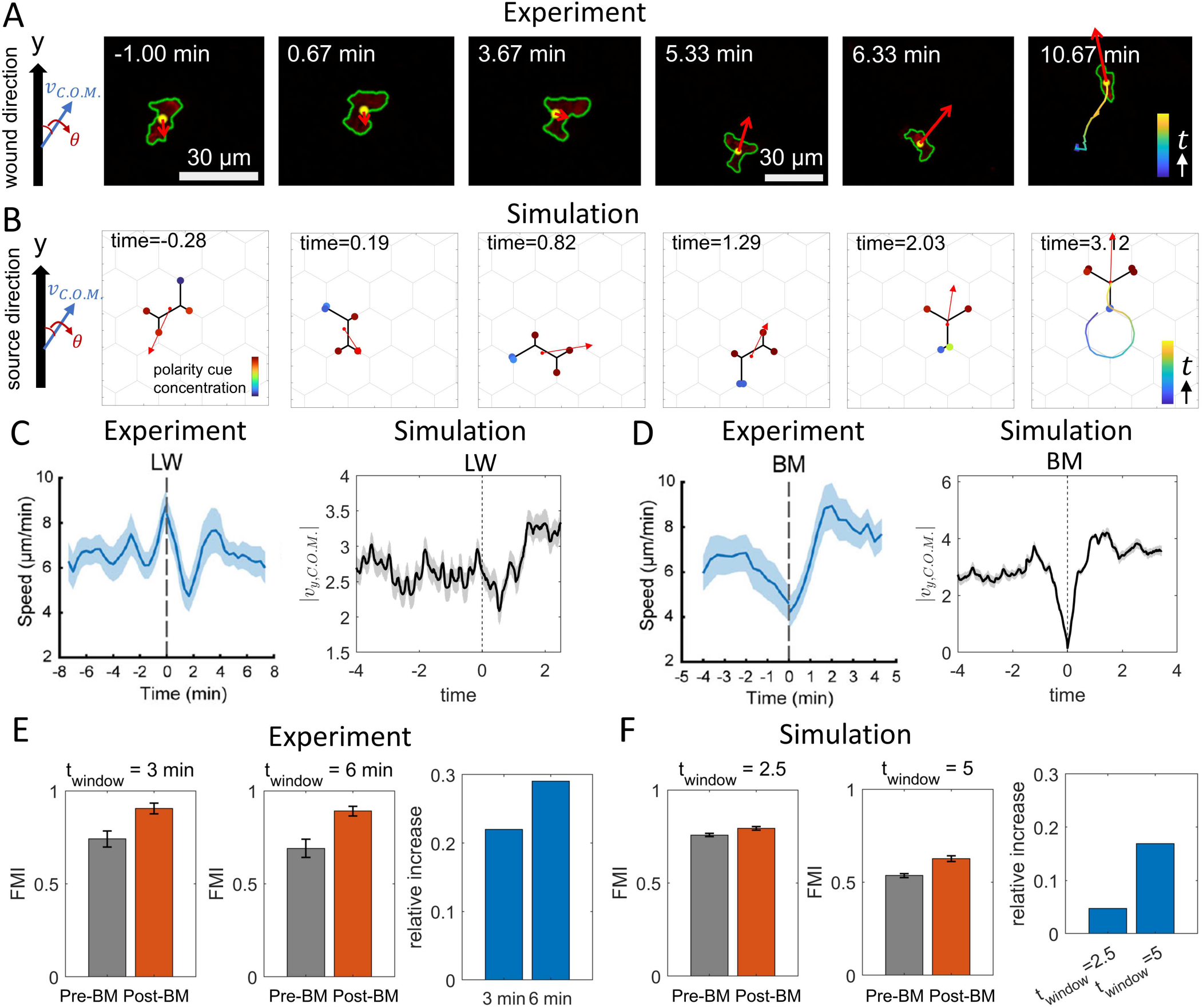
Comparisons between the migration dynamics in simulations and neutrophils in zebra-fish larva [4].(A-B) Snapshots of experimental and simulated cell migration processes. We pick examples where the cell was moving roughly away from the signal, which is located at the positive *y*-direction (large arrow in leftmost panels). At time *t* = 0 the laser-induced wound is applied in the experiments, and the gradient in chemokine is introduced in the simulations. At the final time panel we show the trajectory of the cell’s center of mass (C.O.M.). Red arrows show the instantaneous velocity vector. (C) Left: Experimental cell’s C.O.M. velocity before and after the LW event. Data is obtained from 21 cells. Right: Simulated cell’s center-of-mass velocity in the *y*-direction, |*v*_*y*,C.O.M_.|, before and after the LW event. Data is obtained from 60 simulation runs. (D) Left: Experimental cell’s center-of-mass velocity before and after the time of the beginning of movement (BM). Data is obtained from 18 cells. Right: Simulated cell’s center-of-mass velocity in the *y*-direction, |*v*_*y*,C.O.M._|, before and after the BM time. Data is obtained from 60 simulation runs. (E) FMI of the experimental cell’s center of mass (C.O.M.) within a specific time window (left two panels) and the relative increase after the BM time compared to before the BM time. (F) FMI of the simulated cell’s C.O.M. within a specific time window (left two panels) and the relative increase after the BM time compared to before the BM time (rightmost panel). Key parameters: *ϵ* = 0.1, *d* = 3, *β*_0_ = 12, σ = 0.5.

In Fig. 4C,D we compare the dynamics of the cells in experiments and simulations, when averaged over many trajectories (21 and 18 cells for Fig. 4C and D in the experiments [4]), and 60 simulations). In Fig. 4C, the average cell speed in experiments and simulations are plotted with respect to the LW time. In the simlation, the speed towards the chemokine source in the *y*-direction is used to eliminate the oscillations in the perpendicular direction (*x*-direction) caused by the hexagonal grid topology, which induces significant fluctuations in the overall velocity. In Fig. 4D, the overall experimental speed and the *y*-direction simulation speed are plotted with respect to the time where the cell was first observed to start moving towards the source (denoted as “beginning of movement” BM time, see [4] and SI Section S-6). In Fig. 4C,D the transient slowing down after the chemokine gradient is introduced is clearly observed, in both experiments and simulations. Overall, the simulations are found to reproduce the qualitative features of how neutrophils respond to the chemokine source, and provide a mechanistic explanation for the origin of these dynamics.

Let us next note the mapping of length and time scales between the experiments and model. In the experiments the cells were ∼ 20 *µm* in length (Fig. 4A) and moved at speed of ∼ 6 *µm*/min on average (Fig. 4C). The simulated cell (for *β*_0_ = 12) has an average length 9.38 and speed of 3.59, both in simulation units. This means that the experimental cell covers about 1 cell length in a 3 min interval. This corresponds to choosing a simulation time interval of ∼ 2.5, such that the simulated cell also covers a distance which corresponds to about 1 cell length. Furthermore, to facilitate the comparison between the theoretical model and the experiments, we present all the data as dimensionless or normalized quantities.

In Fig. 4E,F, we compare the effects of the chemokine presence on the FMI of the neutrophils [4] with our simulation model. We use this experimental data to calibrate the value of the parameter *β*_0_ in our simulations. The FMI of the wild-type cells is observed to significantly increase after the BM time (by ∼ 20% − 30%, Fig. 4E). This behavior is captured by the model only when we are in the regime of large *β*_0_.

We next compared the model predictions to the observations of chemotaxis on cells treated with various drugs that inhibit the cytoskeletal activity. We incorporate the inhibition of actin polymerization or myosin-II activity in our model through changes to the model parameters, as was done previously [10]. Comparing the model to the experiments on these drug-treated cells (Fig. S-7) gives good qualitative agreement, and points to the WT cells residing in the high-*β*_0_ regime of our model.

Our conclusion from the comparisons above, that the neutrophils correspond to the high-activity cells of our model, is further supported by the following features of their migration. When the neutrophils start their trajectory further away from the LW, we often find that cells exhibit highly meandering paths (Fig. 5A(i)). Such meandering paths are found in our model only for the high-*β*_0_ regime (see Fig. 3D). Furthermore, we can often identify that the cells display the slow-mode behavior along these trajectories (Fig. 2C(iii)), as shown in Fig. 5A(ii,iii) (the prevalence of these events is quantified in Fig. S-8). Very similar features of cell migration and shape dynamics are observed for a PLB-985 (promyelocytic leukemia blasts) cell migrating *in-vitro* towards a linear chemokine gradient (Fig. 5B), while confined within a regular hexagonal lattice of pillars (see SI section S-2 for more details about these experiments [9]). This demonstrates that the migration characteristics are not dependent on specific *in-vivo* interactions of the neutrophils with the surrounding tissue cells, but are general features of the chemotactic migration of fast-moving branched cells within a complex environment. A simulation of a high-activity cell migrating towards the LW predicts meandering paths with slow-mode events (Fig. 5C). Note that the total cell length increases in the slow-mode events (both in experiments and simulations, Fig. 5A-C panels (ii)) as the two arms extend, before one of them retracts and allows the cell to resume its migration. Note that in the simulations the slow-mode events tend to be more persistent as the cell is closer to the chemokine source, due to the increased actin activity (Figs. 5C(ii),S-8).

**Fig. 5:**
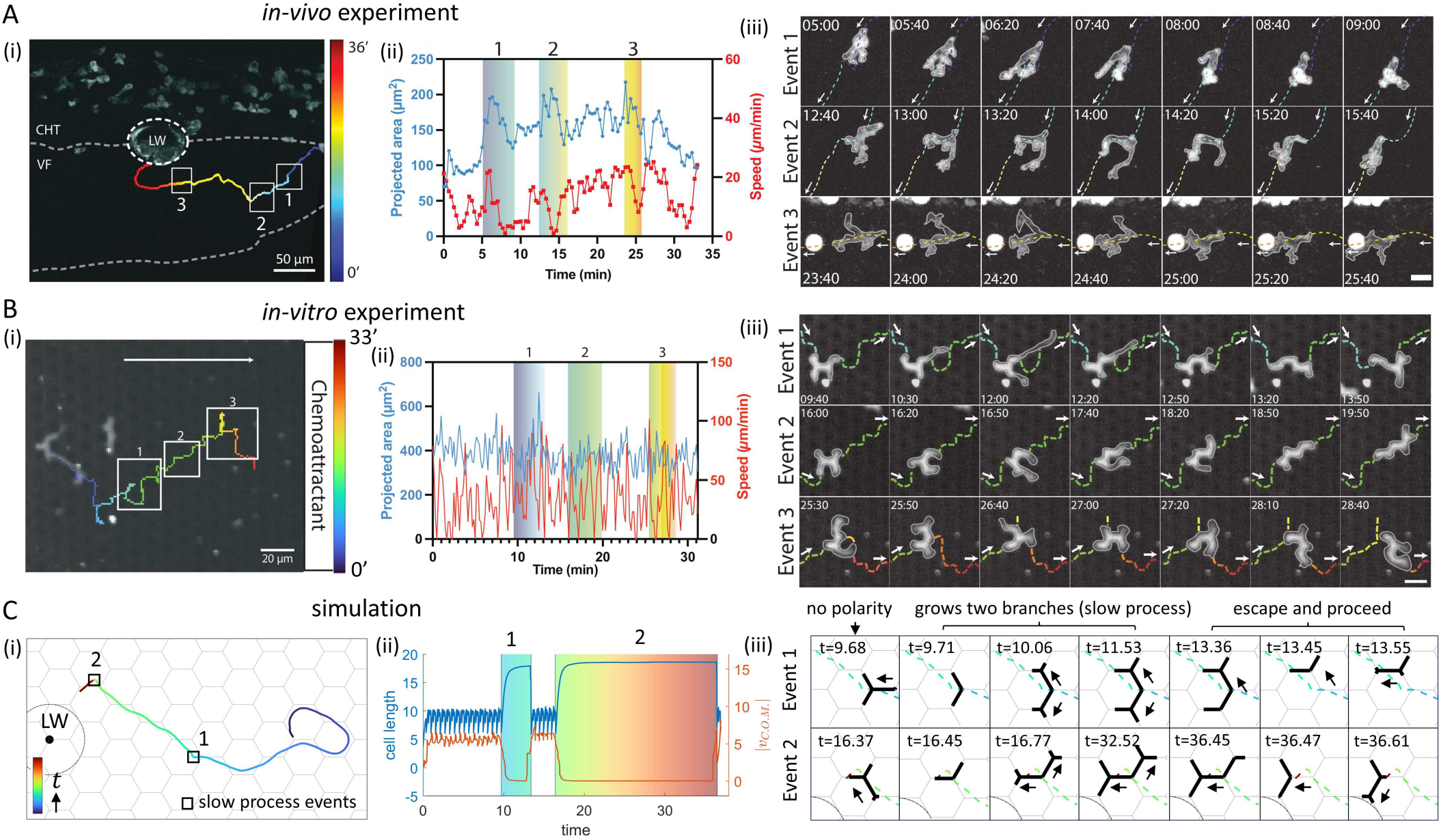
Comparisons between the migration dynamics in simulations and neutrophils. (A) (i) A representative trajectory of a neutrophil migrating toward the LW, with the trajectory color indicating migration time. White boxes highlight time periods along the trajectory in which slow-mode events occur, corresponding to the images in (iii). (ii) Dynamics of the projected area and the overall speed of the neutrophil during migration. The three colored regions correspond to the three slow-mode events marked in (i). (iii) Snapshots of the neutrophil during the three slow-mode events. (B) Same panels as in (A) for a PLB-985 (promyelocytic leukemia blasts) cell migrating *in-vitro* up a linear chemokine gradient, while confined within a regular lattice of pillars. (C) (i) A representative simulation trajectory of a cell migrating toward the LW, with the trajectory color indicating migration time. Black boxes highlight time periods along the trajectory in which slow-mode events occur, corresponding to the color regions in (ii) and the images in (iii). (ii) Dynamics of cell length and its C.O.M. speed during migration. The two colored regions correspond to the three slow-mode events marked in (i). (F) Simulation snapshots of the cell during the two slow-mode events (marked by boxes in (i)). Parameters: *C/c*_0_ = 0.01, *ϵ* = 0.1, *d* = 3.7, *β*_0_ = 12.6, σ = 0.6.

## CONCLUSION

We presented here a theoretical model of the chemotaxis of branched cells on networks. The model exhibits a speed-accuracy trade-off, and predicts that fast cells are less accurate than slow cells. The model provides a mechanistic description of how motile cells respond, reorient and migrate towards a chemoattractant source, in good agreement with experimental observations of immune cells *in-vivo*.

Comparing the model to experimental observations of neutrophils migrating *in-vivo* to the site of a laser-inflicted wound in a zebrafish larva fin, and to PLB-985 (promyelocytic leukemia blasts) cells migrating *in-vitro* within a regular lattice of pillars, suggests that these immune cells correspond to the fast-cell limit of the theoretical model. Cells in this limit have the advantage of minimizing their arrival time to the wound site, if they start close to the wound and receive a strong chemokine signal. This can therefore be a good strategy if the neutrophils are uniformly spread at sufficiently high concentration in the skin, so that there are always some cells close to any wound site. However, such fast moving cells perform less accurate chemotaxis when further from the wound, where the original wound-secreted signal is weak. For efficient recruitment of these far-field neutrophils the immune cells need to utilize different mechanisms, such as secretion of their own chemokine signals. Indeed, neutrophils have evolved such recruitment mechanisms, which have been discovered and are being studied [19–22].

We demonstrated here that our simplified theoretical framework gives a description of both the microscopic dynamics of cell polarization and shape changes, as well as the large-scale migration patterns. This can be a useful tool for deciphering the behavior of motile cells, such as immune or cancer cells, performing chemotaxis inside the complex geometries of living tissues.

## ACKNOWLEDGEMENTS

N.S.G. is the incumbent of the Lee and William Abramowitz Professorial Chair of Biophysics (Weizmann Institute), and acknowledges support from the Royal Society Wolfson Visiting Fellowship, and Human Frontier Science Program grant RGP0032/2022. Work by M.S., I.W., G.R. and A.G. was supported by the Leverhulme Trust (grant RPG-2021-226) and the European Research Council (ERC) under the Horizon 2020 program and UKRI, Grant agreement No. EP/Y02799X/1. M.S. and I.d.V acknowledge support by the European Research Council (grant ERC-SyG 101071793 to M.S), and thank Jack Merrin for support with microfluidic engineering.

