## Supplementary Information for "Chemotaxis of branched cells in complex environments"

### Supplementary Information - Chemotaxis of branched cells in complex environments

Jiayi Liu[1,2], Jonathan E. Ron[2], Giulia Rinaldi[3], Ivanna Williantarra[3], Antonios Georgantzoglou[3,4], Ingrid de Vries[5], Michael Sixt[5], Milka Sarris[3] and Nir S. Gov[2,3]
[1]*Department of Physics, Yale University, New Haven, CT, USA*
[2]*Department of Chemical and Biological Physics, Weizmann Institute of Science, Rehovot, Israel*
[3]*Department of Physiology, Development and Neuroscience, Downing Site, University of Cambridge, Cambridge, UK*
[4]*Novo Nordisk Foundation Center for Stem Cell Medicine (reNEW), Department of Biomedical Sciences, University of Copenhagen, Denmark and*
[5]*Institute of Science and Technology Austria (ISTA), Klosterneuburg, Austria*

#### S-1. *In-vivo* experimental methods

##### Zebrafish Husbandry and Preparation

Tg(lyzC:Gcamp6f) transgenic zebrafish line were maintained adhering to the UK Home Office regulations, UK Animals (Scientific Procedures) Act 1986, which was reviewed by the University Biomedical Service Committee. Adult zebrafish were maintained and bred according to standard protocols [1]. Embryos were collected at 3 hours post fertilisation, bleached for 5 minutes using 0.003% NaOCl (Cleanline, CL3013), followed by three times washing in E3 media (as described in [2]). Embryos were then raised at $28^oC$ in E3 medium supplemented with 0.003% PTU (1-phenyl 2-thiourea; Sigma Aldrich, P7629) to inhibit pigmentation and maintain optical transparency. 3 days post-fertilization (dpf) larvae were used on the day of imaging.

##### Bacterial Culture and Preparation

A day before the imaging day, a single colony of Pseudomonas aeruginosa (strain PAO1 from Martin Welch), isolated via a four-way streak on Pseudomonas Isolation Agar (BD Difco, 292710) supplemented with cetrimide and nalidixic acid (E&O Laboratories Ltd, LS0006) and glycerol (Fisher Scientific, G/0650/17), was grown in 5 mL of antibiotic-free LB broth (Formedium, LBX0102) at $37^oC$ with shaking for 24 hours. On the imaging day, the overnight culture was diluted and incubated to logarithmic phase ($OD_{600}$ = 0.6–0.8). Bacterial concentration was adjusted to $310^5$ CFU/ml using spectrophotometric estimation, based on the assumption that $OD_{600} = 0.5$ corresponds to $10^7$ CFU/ml (based on our unpublished data). Bacteria were pelleted and washed twice with PBS (Sigma Aldrich, D8537) to remove residual medium before resuspension in Ringer solution (as described in [3]) containing 0.16 mg/ml (1x) Tricaine (Merck, A5040).

##### Zebrafish Mounting and Infection

Transgenic larvae expressing the most prominent calcium indicator, Gcamp6f (cDNA originally described by [4]), were screened to ensure high expression. Larvae displaying strong fluorescence and healthy morphology were selected for imaging. Anesthetised larvae were mounted in a 1:1 mix of 2% low-melting agarose (Invitrogen, 16520) and 2× Tricaine in a custom-built coverslip chamber, composed of glass coverslips covering both the top and bottom of the samples and sealed onto a metallic ring. Larvae were oriented laterally and agarose was allowed to solidify. After the agarose was solidified, the tail fin was exposed by carefully removing the surrounding agarose under a dissecting microscope using a tweezer and a capillary needle, allowing bacterial access post-wounding. The chamber was then filled with 1 ml of Ringer solution containing 1x Tricaine and the diluted PAO1 suspension before sealing.

##### Two-Photon Imaging and Laser Wounding

Laser ablation [3] and time-lapse imaging were performed using a LaVision TriM Scope multiphoton microscope equipped with an electro-optic modulator for rapid power modulation. A Spectra-Physics Insight DeepSee dual-line laser was tuned to 900 nm for imaging and 1,040 nm for ablation, with the imaging power set to $\sim 500$ mW at the specimen plane. Image acquisition was performed using ImSpector Pro software (5.0.284.0, LaVision Biotec, ©1998-2016). Two-photon microscopy was performed using a 25×/NA 1.05 water dipping objective lens, with $ddH_2O$ applied to the lens and the correction collar set to 0.17 to match the refractive index. Out of all the mounted larvae, the larvae with the best overall health (normal circulation, heartbeat and morphology) and no damages or aberrant wounding due to mounting was selected for image acquisition. The ventral fin within the caudal hematopoietic tissue of the selected larvae was then centered using brightfield optics. A region of interest with a diameter of 40 $\mu$m was defined as the wound area on a single focal plane (as superficial as possible), scanned at 240 nm/pixel with a 15 $\mu$s dwell time. Z-stacks were acquired every 20 s with a step size of 2 $\mu$m, consisting of 20 planes. Laser wound was triggered after a 2-minute pre-wound baseline, followed by 3 hours of post-wound imaging. Focus was maintained throughout acquisition. Image stacks were processed in Fiji (ImageJ 1.52p, June 2019, publicly available; [5]) using maximum intensity z-projections to generate neutrophil swarming timelapse videos of which were then used for subsequent analysis.

##### Imaging Analyses

Analysis of neutrophil trajectories was performed in Imaris v8.2 (Bitplane AG) on 2D maximum intensity projections of the 4D time-lapse movies (*in-vivo* experiments) and 3D time-lapse movies (*in-vitro* experiments) (as previously described in [2, 6]). Cell trajectories were manually tracked overtime, speed and straightness coefficient of the trajectories were calculated with the imaging software. The projected area of the cells overtime was calculated by manually drawing the perimeter of the cells in Fiji (Image J) and using the measure tool.

#### S-2. *In-vitro* experimental methods for cells migrating in a regular array of pillars

We describe here the microfabrication process of the PDMS devices, their design and manufacturing (Fig.S-1) [7].

##### Photomask

The Photomask is chrome patterned on a glass plate and used to fabricate the wafer. Photomask design is drawn in CorelDraw X8 and exported to dxf file format. LinkCad is used to convert dxf to Gerber format. Photomasks are manufactured by https://www.jd-photodata.co.uk/

##### Wafer

The wafer was backed for 5 min at 110$^o$C, followed by a spin coat Su8 6005 TF (MICROCHEM) and pre-backed for 5-10 at 110$^o$C. The wafer was exposed to 100 mJ/cm$^2$ and post backed at 110$^o$C for 5 min. This was followed by a developing step with SU8 developer and IPA and 135$^o$C baking for 5 min. The height of the device was measured on a profilometer. Finally the wafer was salinized with 1H,1H,2H,2H perfluorooctyltrichlorosilane for 1 h in a sealed vacuum.

##### PDMS devices

40 ml 1:10 PDMS Sylgard 184 (Dow Corning) was mixed and degassed for 2 min at 2000 rpm and for 2 min at 2200 rpm in a Thinky ARE-250 mixer/defoamer and poured on the wafer in an aluminum mold, degassed in a desiccator and cured for 2 h at 80$^o$C. The devices were cut in small squares and 2,5-mm holes were punched (harris Unicore biopsy puncher) on both sides as loading ports. The devices were cleaned with tape (scotch magic tape) and air blown. Cover slip (#1,5, 22x22, Menzel) and device were plasma activated for 2 min in a plasma cleaner (Harrick Plasma). The activated side of the device was placed on the charged side of the coverslip and baked at 95$^o$C for 15 min to achieve bonding. Devices attached to the coverslips were glued with paraplast x-tra (sigma) onto the bottom of a 6 cm cell culture dish so that it covers a central hole of 17 mm diameter. 1-2 h before adding cells devices were incubated with R10 media (RPMI 1640 media (21875091 Gibco) supplemented with 10% FCS (Gibco) and Penicillin-Streptomycin), in a cell culture incubator at 37$^o$C and 5% $CO_2$.

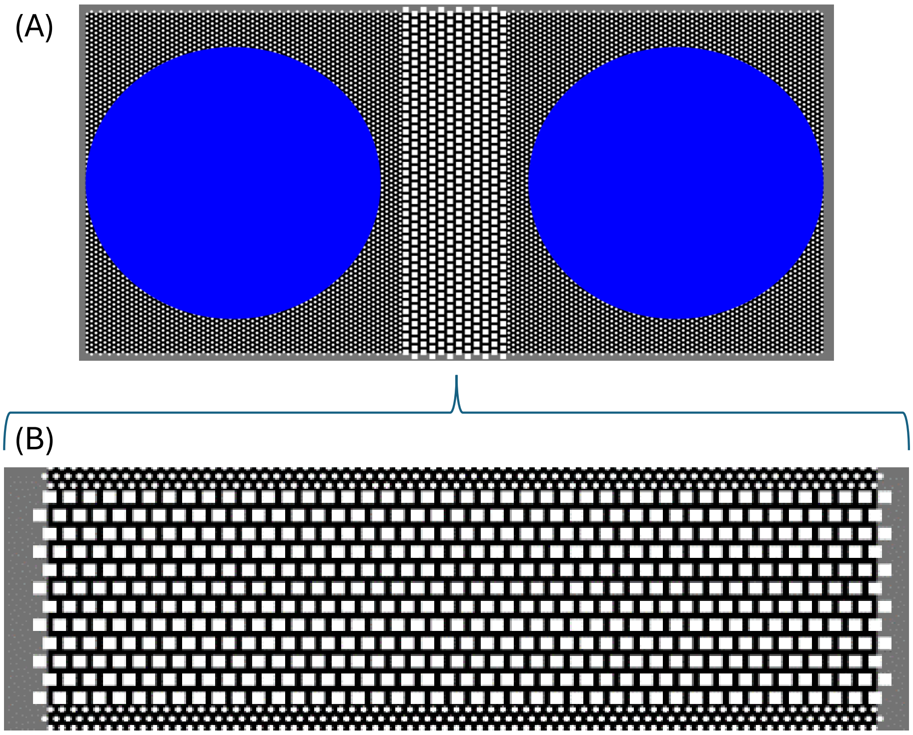


Fig. S-1: Microfabricated PDMS devices design. (A) Blue dots indicate loading areas for cells and chemoattractant. Loading areas are surrounded by an area of circular pillars, which serve as an antechamber from where cells enter the 0,7 x 2,5mm analysis area. (B) Analysis area harbors 10x10$\mu$m rectangular pillars of 3.8$\mu$m height. Pillars are distanced 5$\mu$m and 3$\mu$m in vertical vs. horizontal direction.

**PLB-985 (promyelocytic leukemia blasts) cells**

PLB-985 cells (promyelocytic leukemia blasts) were obtained from the DSMZ (PLB-985 ACC 139). Cells were grown in R10 media. 3-4 days before the experiment, cells where differentiated by adding 1,25% of DMSO to the R10 media. Before the experiment cells were stained with 10$\mu$M TAMRA (Invitrogen) diluted in PBS for 5 min in the dark and washed 3x in R10 media.

Imaging cells in micropillar devices Liquid was removed from both loading ports. One port was filled with 5-7$\mu$l of fMLP (50 mM) and the other port with 5-7$\mu$l of cell suspension (20.000 cells/$\mu$l). The loaded devices were placed in the incubator for minimum 1 h. Movies of cells migrating in the devices were acquired with an imaging interval of 10 s using a Nikon Ti2E inverted widefield Microscope equipped with a Plan Apo λ 20x/0.75 DIC 1 air PFS objective, a monochrome CMOS sensor camera and a custom-built climate chamber (37$^o$C, 5% CO2, humidified).

#### S-3. Model equations

For a cell with $N$ ($N \geq 3$) arms, i.e., spanning across $N-2$ junctions (Fig. S-2), the dynamics of the arm $i$ are described by three variables: 1) its length $x_i$, 2) the fraction of active slip-bonds adhesion $n_i$ at the leading edge, and 3) the local actin treadmilling flow velocity $v_i$ at the leading edge. The dynamic equations for these variables are given by [8]

$$\dot{x}_i = \frac{1}{\Gamma_i}\left[v_i - k\,(L-1)\right] \tag{S-1}$$

$$\dot{n}_i = r\,(1-n_i) - n_i\,\exp\left[\frac{-v_i + k\,(L-1)}{f_s\,n_i}\right] \tag{S-2}$$

$$\dot{v}_i = -\delta\left(v_i - v_i^*\right) + \sigma\xi_t \tag{S-3}$$

In Eq.S-1, $\Gamma_i$ is a non-constant friction coefficient that depends on the direction of motion of the arm's leading edge, given by

$$\Gamma_i = \Theta(\dot{x}_i) + (1 - \Theta(\dot{x}_i))\, n_i \, \exp\left[\frac{v_i - k\,(L-1)}{f_s\, n_i}\right] \tag{S-4}$$

where $\Theta$ is a Heaviside function and $\kappa$ is the effective spring constant of the bond-linkers. When the arm is extending, the friction acts as a constant drag $\Gamma_i = 1$, while when the arm is retracting, the friction is due to the adhesion of the slip-bonds.

Eq.S-1 is a simplified description of the protrusive traction forces, which can be further elaborated to include the adhesion dependence of these forces [9]. The restoring force of the cell elasticity in Eqs.S-1,S-2 is described by a simple spring term, where $k$ is the effective elasticity of the cell (of rest length 1).

In Eq.S-2, $f_s$ describes the susceptibility of the slip-bonds to detach due to the applied force, and $r$ is the effective cell-substrate adhesiveness.

In Eq.S-3, $\delta$ is the rate at which the local actin flows relax to the steady-state solutions $v_i^*$, which is given by

$$v_i^* = \beta \frac{c_s}{c_s + c_i(x_i)} \tag{S-5}$$

where $c_i(x_i)$ is the concentration of actin polymerization inhibitor at the tip of arm $i$ (at the coordinate $x_i$), $c_s$ is the saturation concentration, and $\beta$ is the maximal actin polymerization speed at the arm edges. The term $\sigma\xi_t$ in Eq.S-3 describes the noise in the actin polymerization activity, with a random Gaussian form of amplitude $\sigma$.

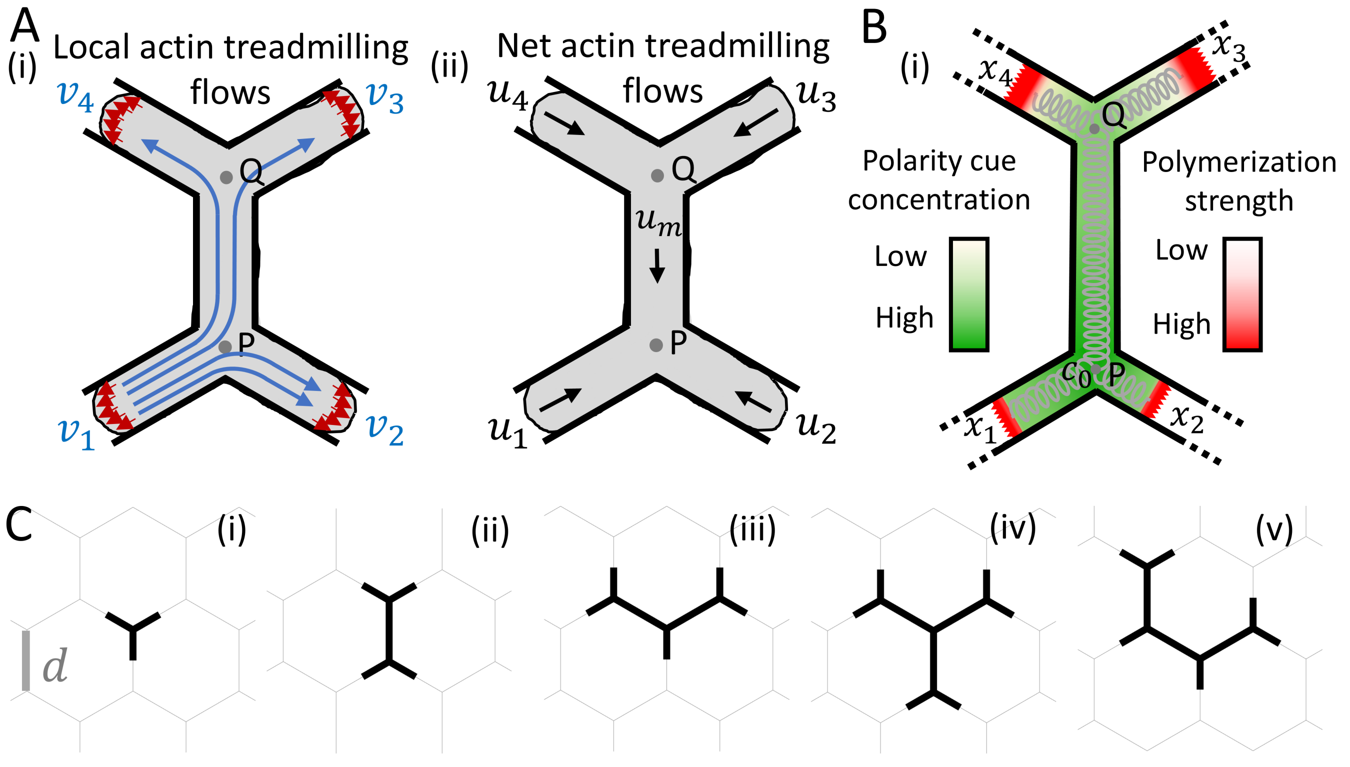


Fig. S-2: (A) (i) Illustrations of the local actin treadmilling flows at each edge of the arms, and the split of the local actin flow of arm 1. (ii) Illustrations of the net actin treadmilling flows in each segment of the cell. (B) An example of the concentration field of the polarity cue, which is affected by the advection flows. The cell elasticity is denoted by the grey springs. (C) Shape of the cells that are spanning different number of junctions. (i) One, (ii) two, and (iii) three junctions. (iv-v) Two possible cell shapes for cells spanning four junctions.

The calculation of the spatial distribution of the inhibitor concentration along the different branches $c_i(x_i)$ was performed. The spatial distribution is composed of exponential sections, maintaining continuity at the junctions, a no-flux boundary condition, and a constant total amount of inhibitor within the cell.

The total length of the cell is given by

$$L = \sum_i x_i + (N-3)\, d \tag{S-6}$$

where $d$ is the distance between two adjacent junctions on the network. Following the initialization of $x_i$, $n_i$ and $v_i$ of each arm, their temporal evolution during the migration process can be obtained through numerical integration of Eq.S-1, Eq.S-2 and Eq.S-3. In this study, we employed the symmetric initial condition.

| Parameter | Value |
|---|---|
| $c$ | 3.85 |
| $D$ | 3.85 |
| $k$ | 0.8 |
| $f_s$ | 5 |
| $r$ | 5 |
| $\kappa$ | 20 |
| $\delta$ | 250 |

Table S-1: Some of the model parameters used in this study.

#### S-4. Effect of noise and hexagon size on chemotactic migration

We investigated the effect of cellular internal noise on the efficiency of chemotactic migration. As shown in Fig. S-4A-C, for both the strong and weak signal regimes (Fig. S-4A and B, respectively), an increase in noise leads to an increase in arrival time and path length, as well as a decrease in $y$-direction velocity and Forward Migration Index (FMI), indicating a reduction in migration efficiency.

In Fig. S-4B, for large $\beta_0$, we see a saturation of the mean path length. This might indicate that the accuracy of cellular DDM has a saturation, such that for $\beta_0 > 8$ there is no significant further increase in decision-making errors. This is also consistent with the mean arrival time which continues to decrease as $\beta_0$ increases, since the path length remains similar, but the migration speed increases with $\beta_0$.

In Fig. S-4C, we calculated the probability of the cell arriving at the target within the maximal simulation time $T = 1000$, $P(arrival)$, as the function of $\beta_0$, for the weak signal regime. It can be observed that for larger noises, there are peaks in $P(arrival)$ at the minima of $\langle T_{arr} \rangle$ from Fig. S-4B(i), while the remaining feature is the minimum in $P(arrival)$ around $\beta_0 \sim 8$.

In Fig.S-4D,E, we investigated the dependence of mean $T_{\rm arr}$, $L_{\rm path}$, $v_y$, and FMI on $\beta_0$ for larger grid size $d = 7.5$. For large $\beta_0$, cells exhibit slow-mode events during migration, with their probability, $P(slow)$, increasing significantly as $\beta_0$ increases (Fig. S-4D,E(iv)). This leads to a substantial increase in $T_{\rm arr}$.

In the strong signal regime (Fig. S-4D), when $\beta_0$ reaches the threshold where slow events occur, a further increase in $\beta_0$ results in a significant rise in $T_{\rm arr}$, while $L_{\rm path}/d$ decreases and FMI increases, indicating an improvement in

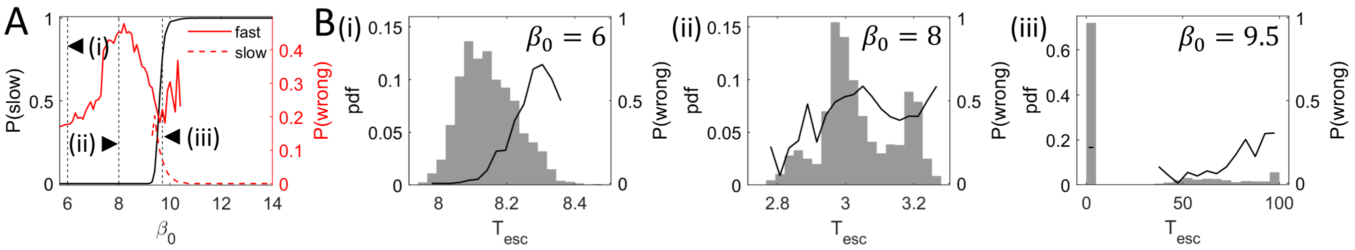


Fig. S-3: (A) For the system in main text Fig.2B(i), we plotted the mean probability of the cell getting into the "slow-mode" on the junction $P(slow)$ (black line). We plotted separately the accuracies of the fast and slow processes, by red solid and dashed lines respectively. (i-iii) Distributions of $T_{esc}$ (histograms) and $P(wrong)$ (solid line) as functions of $T_{esc}$, for the values of $\beta_0$ indicated by the vertical dashed lines in (A). Key parameters: $\epsilon = 0.001$, $\sigma = 0.1$.

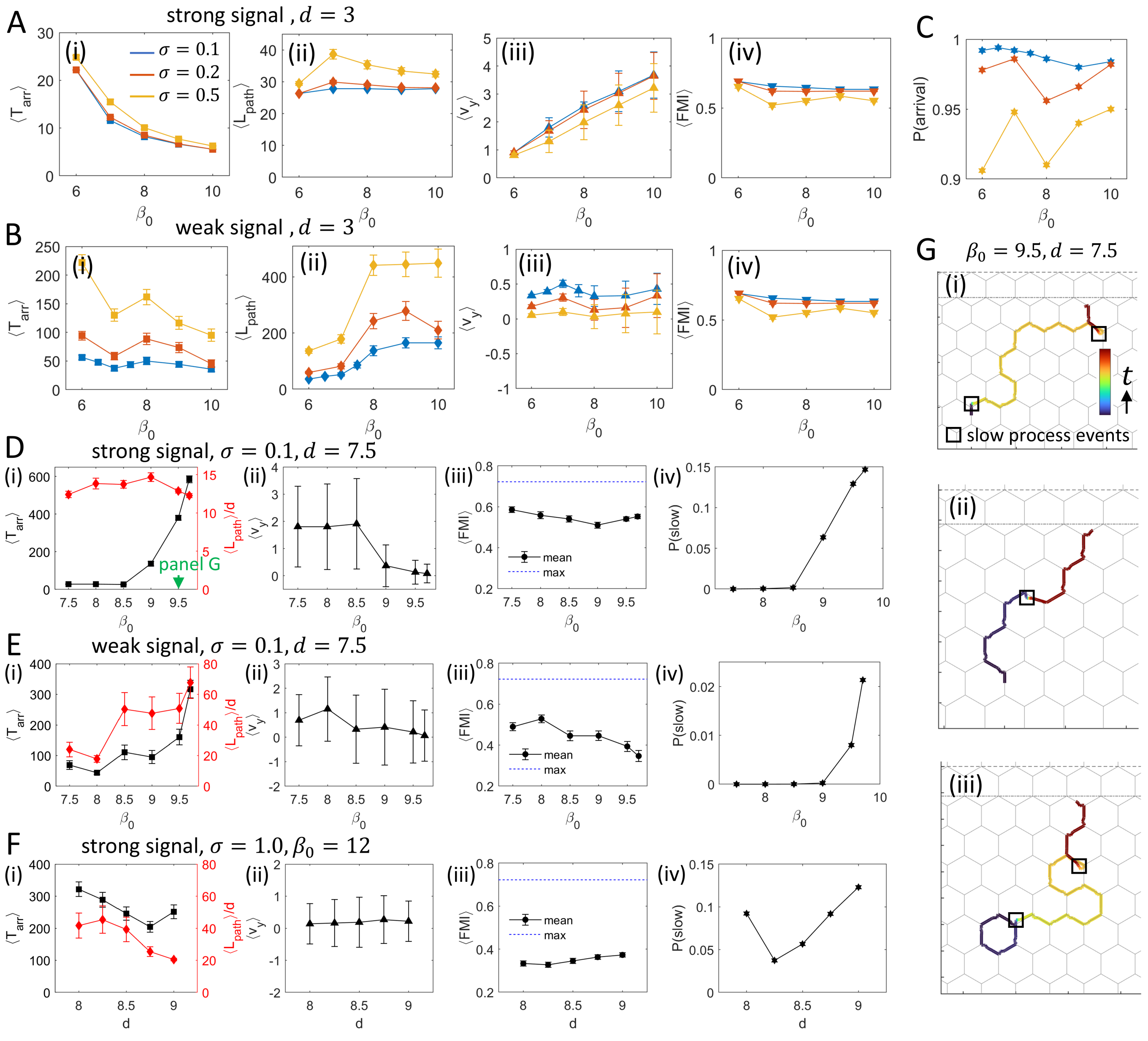


Fig. S-4: Effect of cellular internal noise $\sigma$ and hexagonal edge size $d$ on chemotactic migration. (A-B) On small grids ($d = 3$): (i) $\langle T_{\rm arr}\rangle$, (ii) $\langle L_{\rm path}\rangle$, (iii) $\langle v_y\rangle$, and (iv) $\langle FMI\rangle$ as functions of $\beta_0$. (A) and (B) correspond to the strong signal regime and the weak signal regime, respectively. (C) Probability of arriving at the source at $T_{max} = 1000$, for the weak signal regime of (B). (D-E) On large grids ($d = 7.5$): (i) $\langle T_{\rm arr}\rangle$ and $\langle L_{\rm path}\rangle$, (ii) $\langle v_y\rangle$, (iii) $\langle FMI\rangle$ (black solid line) and the maximal theoretical $FMI$ (blue dashed line), and (iv) $P(\text{slow})$ as functions of $\beta_0$. (D) and (E) correspond to the strong signal regime and the weak signal regime, respectively. (F) Effect of grid size $d$ on: (i) $\langle T_{\rm arr}\rangle$ and $\langle L_{\rm path}\rangle$, (ii) $\langle v_y\rangle$, (iii) $\langle FMI\rangle$, and (iv) $P(\text{slow})$ for $\beta_0 = 12$, in the strong signal regime (as in (A)) and under a large noise level ($\sigma = 1.0$). (G) Typical trajectories of the cell's C.O.M., for $\beta_0 = 9.5$, $d = 7.5$, and $\sigma = 0.1$. Black squares mark the occurrence of slow process events along the trajectory. Maximal simulation time: $T = 1000$. Other key parameter: $\epsilon = 0.2$.

accuracy. In the weak signal regime (Fig. S-4E), the occurrence of slow events also leads to a significant increase in $T_{\rm arr}$, but the increase in $L_{\rm path}/d$ and the decrease in $FMI$ suggest that, in this case, the slow mode does not enhance accuracy.

In Fig. S-4F, we examined the effect of grid size ($d$) on cell migration in the strong signal regime under large $\beta_0 = 12$ and high $\sigma = 1$ conditions. We observed that for larger $d$, the quantities $P_{\rm slow}$ and $FMI$ increase, while $L_{\rm path}/d$ decreases (Fig. S-4F(i,iii,iv)), indicating that the slow mode indeed enhances migration accuracy, consistent with the case of strong signal and small noise (Fig. S-4C). Note that for the cells that do not arrive at the target within the

$T_{\max}$, we assigned $T_{\text{arr}} = 1000$.

In Fig. S-4G we plotted typical trajectories of the cell's C.O.M., for $\beta_0 = 9.5$, $d = 7.5$, and $\sigma = 0.1$. Black squares mark the occurrence of slow process events along the trajectory. Note that these slow-mode events are of a type that is different from those found on a single junction (as shown in Fig.2) and on the hexagonal network in Fig.5F. Instead of the long arms forming in the direction of migration, we find that the cell undergoes a stick-slip event, after which one of the two long arms form along the direction from which the cell arrived at the junction (as shown in Fig.S-9).

#### S-5. Detailed analysis of the dynamics of the cell's trajectory

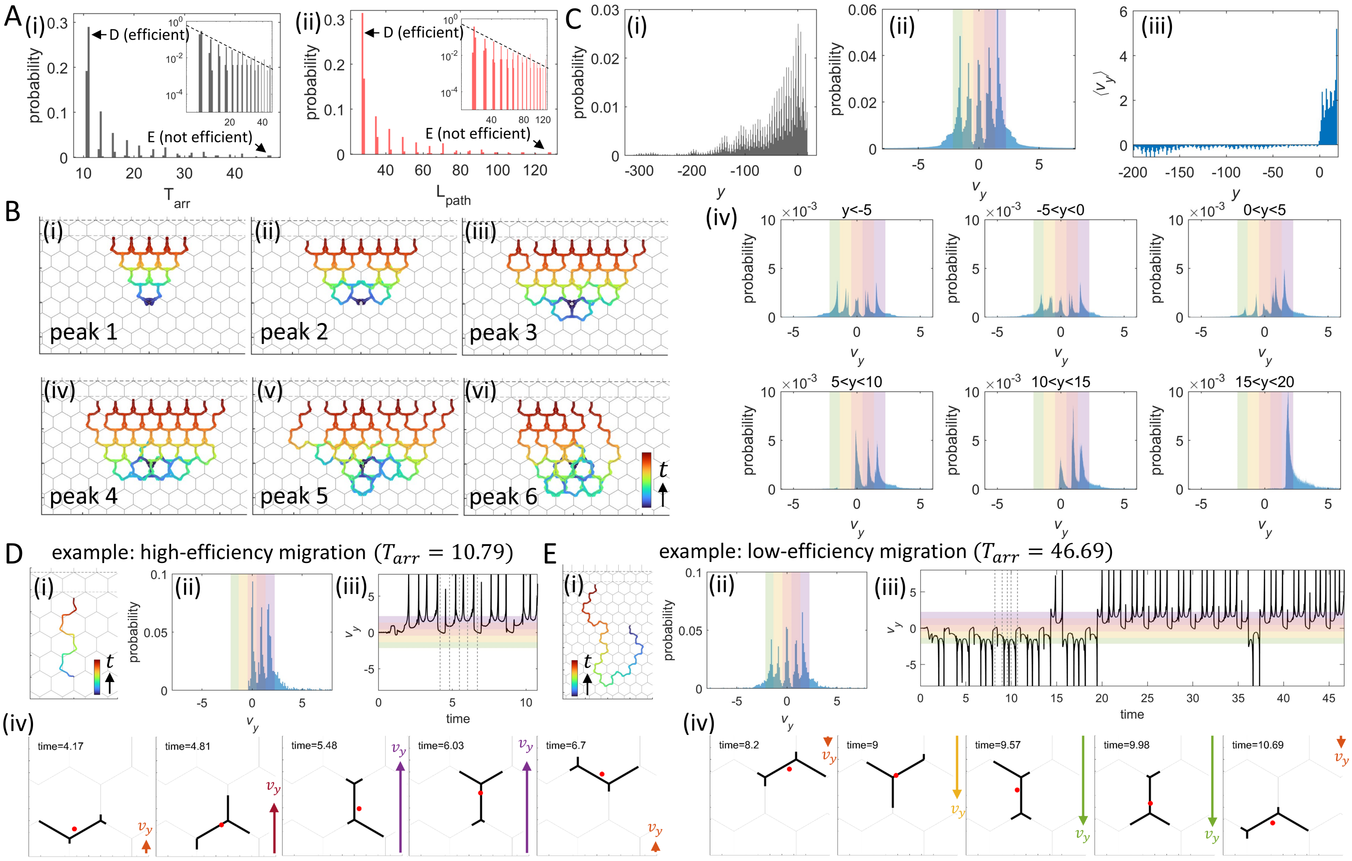


Fig. S-5: Analysis of the $T_{arr}$, $L_{path}$, $y$ and $v_y$ in the slow regime ($C/c_0 = 0.01$). A) Distributions of (i) $T_{arr}$ and (ii) $L_{path}$. B) Trajectories corresponding to the first six peaks of the $T_{arr}$ distribution and the $L_{path}$ distribution in A. C) Distributions of (i) $y$ and (ii) $v_y$ of the cell's centroid. (iii) The $v_y$ distributions at different $y$ ranges. (iv) The $\langle v_y \rangle$ as a function of $y$. D) Example of high efficiency migration. (i) Trajectory of cell's C.O.M.. (ii) Distribution of $v_y$ during the trajectory. (iii) Time series of $v_y$ during the trajectory. (iv) Snapshots for the time stamps (gray dashed lines) in (iii). Key parameters: $\epsilon = 0.2, d = 3, \beta_0 = 8, \sigma = 0.1$.

We investigated the dynamics of the cell's trajectory in more detail for the weak signal regime. We plotted the distributions of $T_{arr}$ and $L_{path}$ (Fig. S-5A), using $\beta = 8$ as an example. Both distributions exhibit discrete peaks, with a constant interval between adjacent peaks. By plotting the trajectories corresponding to the first six peaks of $T_{\text{arr}}$ and $L_{\text{path}}$ (Fig. S-5B) we conclude that the constant interval between the arrival time (and path length) peaks arises from the additional time (or path length) that is added to the trajectory each time the cell makes an additional incorrect decision along the path, i.e. a turn that takes it away from the chemokine source.

We also plotted the distribution of $y$ and $v_y$ (Fig. S-5C(i-ii)), where the peaks result from the periodic migration patterns of the cell as it moves along the hexagonal edges. Furthermore, we plotted $\langle v_y \rangle$ (Fig. S-5C(iii)) and the $v_y$ distributions within specific ranges of $y$ (Fig. S-5C(iv)). As expected, $\langle v_y \rangle$ is positive only at positive $y$-positions, where the cell is sufficiently close to the chemokine source. Further away from the source the cell loses the very weak signal and performs isotropic diffusion.

For illustration, we provide examples of efficient and inefficient migration paths (corresponding to the 1st and 15th peaks in Fig. S-5A), shown in Fig. S-5D,E. Panel (i) displays the trajectory, while (ii) shows the $v_y$ distribution. Panel (iii) presents the time series of $v_y$, with shaded regions corresponding to the respective $v_y$ ranges indicated by the colors in (ii). We found that the $v_y$ time series exhibits a highly periodic pattern, as the cell performs stick-slip migration over the regular hexagonal network. This migration pattern, with fast slip events, is further demonstrated in panels (iv), where we present typical snapshots of the cell shape and C.O.M. at the times indicated by the vertical black dashed lines in panels (iii).

##### S-6. Comparing the experimental response of the cell to the chemokine with the model

To simplify the theoretical model, we assumed a spatially linear variation of the chemokine concentration for the comparison with the experimental data (Fig. S-6A(i)). As in the previous theoretical analysis , the concentration is directed along the $y$-axis. It reaches its maximum value $c_0$ at the source position $y_{\text{source}}$. Now, we assumed that at a specific $y$ position, $y_{\text{end}}$, the concentration drops to zero. The concentration varies linearly between $y_{\text{source}}$ and $y_{\text{end}}$, meaning that the chemokine concentration function $c(y)$ in our model (Eq.3) takes the following linear form:

$$c(y) = c_0 \frac{y - y_{\text{end}}}{y_{\text{source}} - y_{\text{end}}} \tag{S-7}$$

We set $y_{\text{source}} = -y_{\text{end}} = 8d$. At the start of each simulation, the cell is symmetrically positioned at the origin $(x, y) = (0, 0)$. The chemokine profile parameters are set as $\epsilon = 0.1$ and $C/c_0 = 1$.

Under this setting, the normalized chemokine concentration and the enhancement of actin polymerization speed as functions of $y$ are presented in Fig. S-6A(ii). It can be observed that although the chosen parameter $C/c_0 = 1$ corresponds to the weak signal ("slow saturation") regime, the linear variation of $c(y)$ ensures that as soon as the chemical cue is introduced, the cell perceives a strong level of signal, provided it is within the region between $y_{\text{source}}$ and $y_{\text{end}}$ (which is indeed satisfied in our setup).

The "Beginning of movement" (BM) time (Fig. 4D) is defined in our simulations as the time point at which cell movement in the direction of the chemical gradient begins (Fig. S-6B). In [6] this time point was defined similarly, where cell motion was first identified as initiated towards the LW signal. It is determined individually for each cell based on the time point at which the cell initiates movement in the "wound" ($y$) direction.

The effect of the chemokine on the FMI of the cells is given in Fig. 5E for the two durations used in [6]. It is clear that the significant improvement in the FMI due to the external gradient, in these short time windows, only appears for large $\beta_0$, which is the regime that fits the experimental observations for the WT cells.

In Fig. S-6C,D(i), we show the dynamics of the cell's C.O.M. velocity in the $y$-direction, $|v_{y,\text{C.O.M.}}|$, and the angle $\theta$ between the C.O.M. velocity vector and the positive $y$-axis for the experiment and the simulation (see schematic in Fig. 4A,B). In both cases, the cells slow down following the introduction of the chemokine gradient, then re-polarize and increase their speed towards the source.

The origin of deceleration in the model can be demonstrated by plotting the maximal front-back difference in the polarity cue along the cell length (Fig. S-6D(ii)), and by plotting the polarity concentrations at the tips of the cellular arms (Fig. 4B). Prior to the introduction of the chemokine gradient, the polarity cue difference oscillates as the cell periodically changes its length and number of arms spanning the junction while moving on the hexagonal network. At time= 0, the cell was polarized such that its leading edge was pointing away from the source. This means that the back of the cell is exposed at time $t = 0$ to a higher chemokine concentration than the front, and therefore the actin polymerization is more enhanced at the cell's rear than at its front. A transient enhancement of the actin polymerization activity at the side of the cell facing the chemokine gradient was also observed in neutrophils [6]. The difference in polymerization activity at the cells edges due to a different chemokine concentration results in a smaller overall front-back retrograde flow and weaker advection of the polarity cue. The front-back polarity cue difference therefore decreases (Fig. S-6D(ii)), which slows down the cell migration. By reducing the cell polarity the protrusive activity becomes more uniform among all the cellular protrusions, not limited to the cell front, and facilitates the rotation of the cell. As the cell re-establishes its front-back polarity in the direction of the chemokine source (the front-back polarity cue difference recovers its large values, Fig. S-6D(ii)), and the cell resumes its fast migration. Note that the oscillations in the simulated cell speed are much more regular than in the experiment (Fig. S-6C,D(i)), but the angle of migration makes oscillations of similar size in both cases.

In Fig. S-6D(iii), we give the dynamics of the arm tips with the largest (front) and smallest (back) $y$-values from the experiment and the simulation used in Fig. 4. The length of the cell along the $y$-axis is shown in Fig. S-6D(iv),

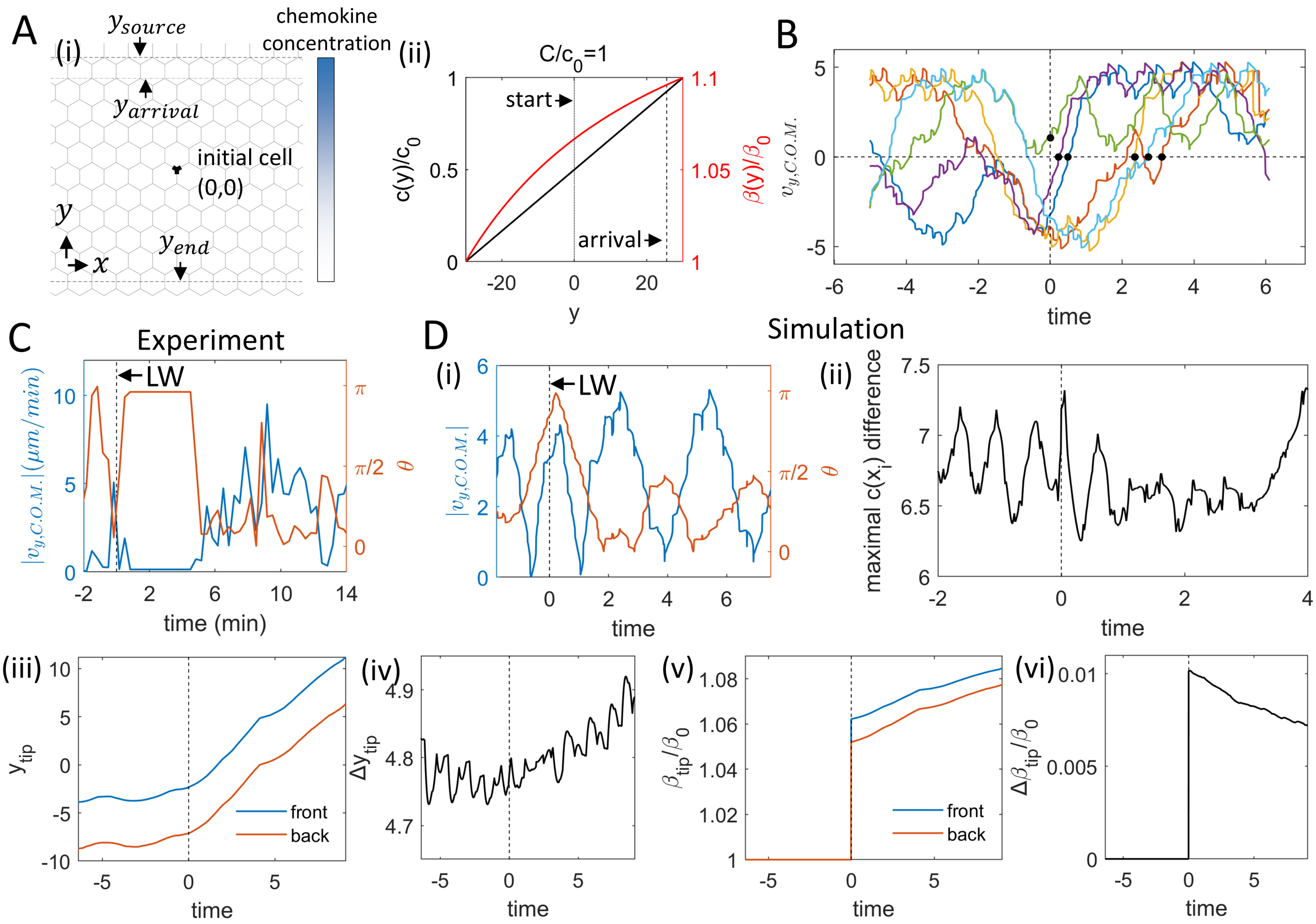


Fig. S-6: Multiple-junction model with a chemokine line source that decays linearly. (A) (i) Schematic representation of the model. Two gray dashed lines indicate the position of the chemokine source and the position where the source decays to zero, respectively. The gray dash-dotted line marks the arrival position. (ii) Normalized chemokine concentration, $c(y)/c_0$ (black line), and the enhancement of actin polymerization speed, $\beta(y)/\beta_0$, as functions of $y$. (B) Dynamics of $v_{y,\mathrm{C.O.M.}}$ for six randomly selected simulation trajectories. The black dots indicate the BM time corresponding to each trajectory. (C) Dynamics of the C.O.M. velocity in $y$-direction, $|v_{y,\mathrm{C.O.M.}}|$, of the experimental cell (main text Fig. 4). (D) Dynamics of the simulated cell's (i) C.O.M. velocity in the $y$-direction, $|v_{y,\mathrm{C.O.M.}}|$, (ii) maximal difference between the polarity cue concentration at the arm tips, $c(x_i)$, (iii) $y$-coordinates of the front tip (maximum $y$-coordinate) and back tip (minimum $y$-coordinate), (iv) difference in $y$-coordinates between the front and back tips, (v) $\beta$ enhancement, $\beta/\beta_0$ at the front and back tips, (vi) difference in $\beta$ enhancement between the front and back tips. Black dashed lines mark the LW time. Parameters: $\epsilon = 0.1, d = 3, \beta_0 = 12, \sigma = 0.5$.

and we find that the cell is stretched by the chemokine gradient in the gradient direction.

The chemokine-induced enhancement of the actin polymerization activity at the front and back tips of the cell (as defined by their $y$-coordinate in (i)), is shown in Fig. S-6D(v). This difference (Fig. S-6D(vi)) is slowly decreasing over time, as the cell moves up the gradient and the overall chemokine concentration around the cell increases.

##### S-7. Comparing the migration characteristics of experimental cells and the model

The total cell length is observed to be significantly diminished by decreasing actin polymerization activity (CK666, Fig.S-7A)[6]. Compared to the WT cells, we fit a reduced $\beta_0$, by $\sim 17\%$ in the CK666-affected cells (Fig. S-7A)[6].

The effect of blebbistatin, which inhibits myosin-II contractility, is described in our model as a decrease in the cell contractile-stiffness parameter $k$ and the retrograde flow of actin $\beta_0$ [10]. While decreasing $k$ acts to make the cell longer, the lower $\beta_0$ decreases the forces that act to elongate the cell, so that combined the cell is similar in length to the WT, as observed in the experiments (Fig. S-7A,B)[6].

In Fig. S-7C,D, we compare the experimental [6] and simulated changes to the FMI for the WT (DMSO) and drug-treated cells. The relative changes are qualitatively captures by the model, especially the increase in FMI due to the chemokine for the WT cells, and the vanishing of this effect upon drug application. This, again, points to the WT cells residing in the high-$\beta_0$ regime of our model, which is the only regime where we see that the FMI is significantly increased by the chemokine gradient.

Similarly in Fig. S-7E,F the normalized cell speed is compared between the WT and drug-treated cells [6]. In the WT cells a large difference is found between the cell speed towards and against the chemokine direction, while this is very much reduced for the drug-treated cells, which also have lower overall migration speed compared to the WT (Fig. S-7E). These qualitative features are captured by the model (Fig. S-7F).

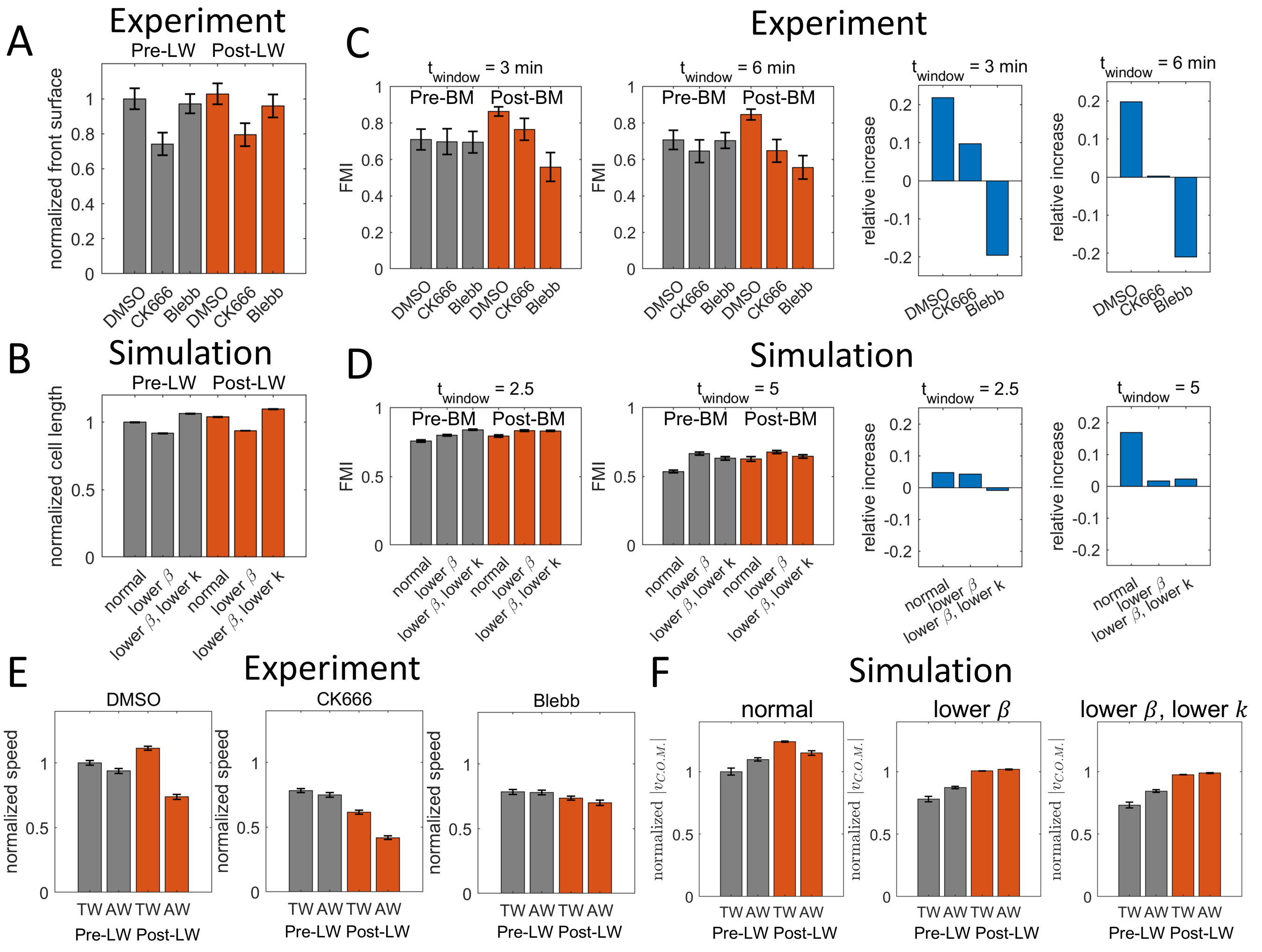


Fig. S-7: Comparison of the model with the chemotaxis properties of Neutrophiles *in-vivo* [6]. (A) Normalized front surface of the experimental cell (normalized by the DMSO value before BM), before and after the BM time (grey and red respectively). (B) Normalized cell length of the simulated cell (normalized by the normal cell value before BM) before and after the BM time (grey and red respectively). (C) FMI of the experimental cell's C.O.M. under different drug treatments within a specific time window (left two panels) and the relative increase after the BM time compared to before the BM time (right two panels). (D) FMI of the simulated cell's C.O.M. under different parameter settings (corresponding to the respective drug treatments in experiments) within a specific time window (left two panels) and the relative increase after the BM time compared to before the BM time (right two panels). (E) Normalized speed of the experimental cell's C.O.M. (normalized by the DMSO value TW(pre)), towards (TW) and away (AW) from the wound, before and after the LW time (grey and red respectively). (F) Normalized speed of the simulated cell's C.O.M. (normalized by the normal cell value TW(pre)), before and after the BM time (grey and red respectively). For wild-type (normal) cells, CK-666-treated cells, and blebbistatin-treated cells, the corresponding simulation parameters were chosen as: $(\beta, k) = (12.0, 0.8)$, $(\beta, k) = (10.0, 0.8)$, and $(\beta, k) = (10.0, 0.7)$, respectively. Other key parameters: $\epsilon = 0.1, d = 3, \sigma = 0.5$.

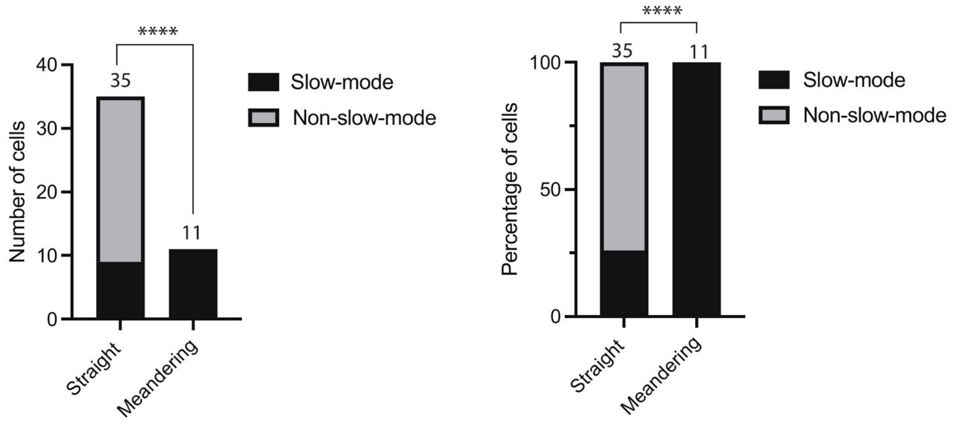


Fig. S-8: Characterizing "slow-mode" events for cells *in-vivo* (as in Fig. 5A-C). GCamp6F transgenic zebrafish embryos (3 dpf) were infected with *Pseudomonas aeruginosa*, laser-wounded, and imaged using a two-photon confocal microscope, as previously described. Neutrophil trajectories were determined manually using the imaging software IMARIS. Meandering and straight trajectories were classified based on the straightness coefficient quantified by the software ($> 0.60$ considered as straight). Trajectories were also categorized based on the presence or absence of slow-mode events. The graph shows both the percentage and absolute count of cells displaying slow-mode events within the straight and meandering categories. Percentages are based on 39 and 11 cells, respectively, as indicated in the graph. Videos with an overall recruitment greater than 15 cells were selected. $N = 46$ neutrophil trajectories were quantified from 7 independent videos/embryos. $****$ $P < 0.0001$, Chi-square test (and Fisher's exact test).


[1] M. Westerfield, *The Zebrafish Book; A guide for the laboratory use of zebrafish (Danio rerio)* (2007).

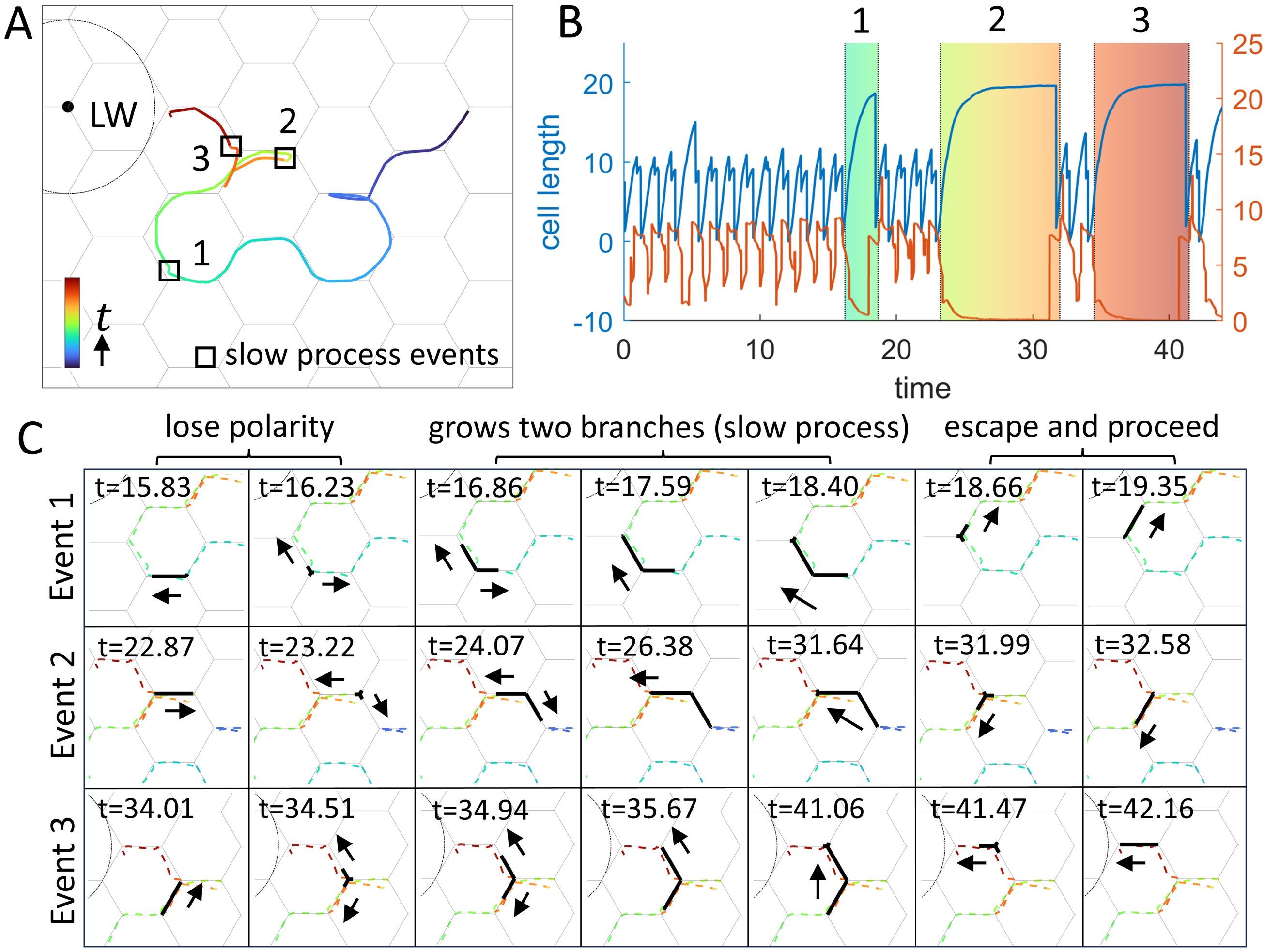


Fig. S-9: Illustration of the second type of slow mode during cell migration on the large grids. (A) A representative simulation trajectory of a cell migrating toward the LW, with the trajectory color indicating migration time. Black boxes highlight time periods along the trajectory in which slow-mode events occur, corresponding to the images in (C). (B) Dynamics of the cell length and the C.O.M. speed of the cell during migration. The two colored regions correspond to the three slow-mode events marked in (D). (C) Simulation snapshots of the cell during the two slow-mode events. Parameters: $C/c_0 = 0.01, \epsilon = 0.2, d = 9, \beta_0 = 12, \sigma = 1.2$.
